# Generating and testing ecological hypotheses at the pondscape with environmental DNA metabarcoding: a case study on a threatened amphibian

**DOI:** 10.1101/278309

**Authors:** Lynsey R. Harper, Lori Lawson Handley, Christoph Hahn, Neil Boonham, Helen C. Rees, Erin Lewis, Ian P. Adams, Peter Brotherton, Susanna Phillips, Bernd Hänfling

## Abstract

Environmental DNA (eDNA) metabarcoding is revolutionising biodiversity monitoring, but has unrealised potential for ecological hypothesis generation and testing. Here, we validate this potential in a large-scale analysis of vertebrate community data generated by eDNA metabarcoding of 532 UK ponds. We test biotic associations between the threatened great crested newt (*Triturus cristatus*) and other vertebrates as well as abiotic factors influencing *T. cristatus* detection at the pondscape. Furthermore, we test the status of *T. cristatus* as an umbrella species for pond conservation by assessing whether vertebrate species richness is greater in ponds with *T. cristatus* and higher *T. cristatus* Habitat Suitability Index (HSI) scores. *T. cristatus* detection was positively correlated with amphibian and waterfowl species richness. Specifically, *T. cristatus* was positively associated with smooth newt (*Lissotriton vulgaris*), common coot (*Fulica atra*), and common moorhen (*Gallinula chloropus*), but negatively associated with common toad (*Bufo bufo*). *T. cristatus* detection did not significantly decrease as fish species richness increased, but negative associations with common carp (*Cyprinus carpio*), three-spined stickleback (*Gasterosteus aculeatus*) and ninespine stickleback (*Pungitius pungitius*) were identified. *T. cristatus* detection was negatively correlated with mammal species richness, and *T. cristatus* was negatively associated with grey squirrel (*Sciurus carolinensis*). *T. cristatus* detection was negatively correlated with larger pond area, presence of inflow, and higher percentage of shading, but positively correlated with HSI score, supporting its application to *T. cristatus* survey. Vertebrate species richness was significantly higher in *T. cristatus* ponds and broadly increased as *T. cristatus* HSI scores increased. We reaffirm reported associations (e.g. *T. cristatus* preference for smaller ponds) but also provide novel insights, including a negative effect of pond inflow on *T. cristatus*. Our findings demonstrate the prospects of eDNA metabarcoding for ecological hypothesis generation and testing at landscape scale, and dramatic enhancement of freshwater conservation, management, monitoring and research.

## 1. Introduction

Environmental DNA (eDNA) analysis offers ecologists exceptional power to detect organisms within and across ecosystems. DNA released by organisms into their environment via secretions, excretions, gametes, blood, or decomposition, can be sampled and analysed using different approaches to reveal the distribution of single or multiple species (Rees et al., 2014; Lawson Handley, 2015). eDNA analysis combined with high-throughput sequencing (i.e. eDNA metabarcoding) can yield efficient, comprehensive assessments of entire communities (Deiner et al., 2017), providing a step change in biodiversity monitoring (Hering et al., 2018). eDNA metabarcoding has untapped potential to generate and test ecological hypotheses by enabling biodiversity monitoring at landscape scale with minimal impact to communities under investigation. This potential has already been demonstrated with targeted eDNA analysis by Wilcox et al. (2018), where climate-mediated responses of bull trout (*Salvelinus confluentus*) to biotic and abiotic factors were revealed using quantitative PCR (qPCR) on crowd-sourced eDNA samples. Although eDNA metabarcoding assessments of alpha and beta diversity along environmental gradients are increasing (e.g. Hänfling et al., 2016; Olds et al., 2016; Kelly et al., 2016; Evans et al., 2017; Li et al., 2018a; Nakagawa et al., 2018), this tool is less commonly used for ecological hypothesis testing, such as the impact of environmental stressors (Li et al., 2018b; Macher et al., 2018).

Aquatic ecosystems are highly suited to eDNA studies as eDNA exists in multiple states with rapid modes of transport and degradation, increasing detectability of contemporary biodiversity (Rees et al., 2014; Barnes & Turner, 2015). Lentic systems provide further opportunities for eDNA research, being discrete water bodies with variable physicochemical properties that do not experience flow dynamics (Harper et al., 2019).Ponds in particular have enormous biodiversity and experimental virtue that has not been maximised in previous eDNA metabarcoding assessments of this habitat (Valentini et al., 2016; Evans et al., 2017; Klymus et al., 2017; Ushio et al., 2017; Bálint et al., 2018). These small and abundant water bodies span broad ecological gradients (De Meester et al., 2005) and comprise pondscapes – a network of ponds and their surrounding terrestrial habitat (Hill et al., 2018). Pondscapes contribute substantially to aquatic and non-aquatic biodiversity across spatial scales, with ponds supporting many rare and protected species in fragmented landscapes (De Meester et al., 2005; Biggs et al., 2016; Hill et al., 2018). Consequently, ponds are model systems for experimental validation and examination of biogeographical patterns (De Meester et al., 2005). Habitat complexity and tools required for different taxa with associated bias (Evans et al., 2017) and cost (Valentini et al., 2016) once hindered exhaustive sampling of pond biodiversity (Hill et al., 2018), but eDNA metabarcoding may overcome these barriers (Harper et al., 2019).

In the UK, the threatened great crested newt (*Triturus cristatus*) is an umbrella species for pond conservation. The extensive literature on *T. cristatus* ecology provides an excellent opportunity to validate ecological patterns revealed by eDNA metabarcoding. Both biotic (e.g. breeding substrate, prey, and predators) and abiotic (e.g. pond area, depth, and temperature) factors are known to influence *T. cristatus* breeding success (Langton, Beckett & Foster, 2001). The *T. cristatus* Habitat Suitability Index (HSI [Oldham et al., 2000; ARG-UK, 2010; O’Brien et al., 2017]) accounts for these factors using 10 suitability indices that are scored and combined to calculate a decimal score between 0 and 1 (where 1 = excellent habitat). Larvae are susceptible to fish and waterfowl predation (Edgar & Bird, 2006; Rannap & Briggs, 2006; Skei et al., 2006; Hartel, Nemes & Oellerer, 2010), and adults reportedly avoid ponds containing three-spined stickleback (*Gasterosteus aculeatus*) (McLee & Scaife, 1992), ninespine stickleback (*Pungitius pungitius*), crucian carp (*Carassius carassius*), and common carp (*Carassius carpio*) (Rannap, Lõhmus & Briggs, 2009a, b). Conversely, *T. cristatus* and smooth newt (*Lissotriton vulgaris*) prefer similar habitat and often co-occur (Rannap & Briggs, 2006; Skei et al., 2006; Rannap et al., 2009a; Denoël et al., 2013; Cayuela et al., 2018). *T. cristatus* individuals thrive in ponds with good water quality as indicated by diverse macroinvertebrate communities (Oldham et al., 2000; Rannap et al., 2009a). Pond networks encourage *T. cristatus* occupancy (Joly et al., 2001; Rannap et al., 2009a; Hartel et al., 2010; Denoël et al., 2013), but larger pond area discourages presence (Joly et al., 2001). Ponds with heavy shading (Vuorio, Heikkinen & Tikkanen, 2013) or dense macrophyte cover (Rannap & Briggs, 2006; Skei et al., 2006; Hartel et al., 2010) are unlikely to support viable populations. *T. cristatus* individuals also depend on terrestrial habitat, preferring open, semi-rural pondscapes (Denoël et al., 2013) containing pasture, extensively grazed and rough grassland, scrub, and coniferous and deciduous woodland (Oldham et al., 2000; Rannap & Briggs, 2006; Rannap et al., 2009a; Gustafson, Malmgren & Mikusiński, 2011; Vuorio et al., 2013).

We assessed vertebrate communities at the pondscape using a dataset produced by eDNA metabarcoding for over 500 ponds with comprehensive environmental metadata. We validated eDNA metabarcoding as a tool for ecological hypothesis generation and testing, and compared its outputs to previous results produced by established methods. Specifically, we identified and tested biotic (community presence-absence data) and abiotic (environmental metadata on ponds and surrounding terrestrial habitat) factors influencing *T. cristatus* detection at the pondscape – an impractical task by conventional means. Furthermore, we tested the applicability of the HSI to predict eDNA-based *T. cristatus* detection. Finally, we assessed the umbrella species status of *T. cristatus* by investigating whether *T. cristatus* detection and the *T. cristatus* HSI score can predict vertebrate species richness of ponds.

## 2. Materials and methods

### 2.1 eDNA samples

Samples were previously collected from 508 ponds as part of Natural England’s Great Crested Newt Evidence Enhancement Programme and from 24 privately surveyed ponds using established methodology (Biggs et al., 2015), detailed in Supporting Information: Appendix 1. Briefly, 20 × 30 mL water samples were collected from each pond and pooled. Six 15 mL subsamples were taken from the pooled sample and each added to 33.5 mL absolute ethanol and 1.5 mL sodium acetate 3 M (pH 5.2). Subsamples were pooled during DNA extraction to produce one eDNA sample per pond. Targeted qPCR detected *T. cristatus* in 265 (49.81%) ponds (Harper et al., 2018).

Environmental metadata were collected for 504 of 532 ponds (Fig. S1) by environmental consultants contracted for Natural England’s Great Crested Newt Evidence Enhancement Programme. Metadata included: maximum depth of ponds; pond circumference; pond width; pond length; pond area; pond density (i.e. number of ponds per km^2^); terrestrial overhang; shading; macrophyte cover; HSI score (Oldham et al., 2000); HSI band (categorical classification of HSI score [ARG-UK, 2010]); pond permanence; water quality; pond substrate; presence of inflow or outflow; presence of pollution; presence of other amphibians, fish and waterfowl; woodland; rough grass; scrub/hedge; ruderals; other terrestrial habitat (i.e. good quality terrestrial habitat that did not conform to aforementioned habitat types); and overall terrestrial habitat quality (see Table S1).

### 2.2. eDNA metabarcoding

We repurposed the taxonomically assigned sequence reads from Harper et al. (2018) that were produced using eDNA metabarcoding of pond water to compare qPCR and eDNA metabarcoding for *T. cristatus* detection. Here, we provide a summary of their eDNA metabarcoding workflow (see Supporting Information: Appendix 1 for details).

A custom, phylogenetically curated reference database of mitochondrial 12S ribosomal RNA (rRNA) sequences for UK fish species was previously constructed for eDNA metabarcoding of lake fish communities (Hänfling et al., 2016). Harper et al. (2018) constructed additional reference databases for UK amphibians, reptiles, birds, and mammals. Reference sequences available for species varied across vertebrate groups: amphibians 100.00% (*N* = 21), reptiles 90.00% (*N* = 20), mammals 83.93% (*N* = 112), and birds 55.88% (*N* = 621). Table S2 lists species without database representation, i.e. no records for any species in a genus. Sanger sequences were obtained from tissue of *T. cristatus, L. vulgaris*, Alpine newt (*Ichthyosaura alpestris*), common toad (*Bufo bufo*), and common frog (*Rana temporaria*) to supplement the amphibian database. The complete reference databases compiled in GenBank format were deposited in a GitHub repository and permanently archived (https://doi.org/10.5281/zenodo.1188710) by Harper et al. (2018). Reference databases were combined for *in silico* validation of published 12S rRNA primers 12S-V5-F (5’-ACTGGGATTAGATACCCC-3’) and 12S-V5-R (5’-TAGAACAGGCTCCTCTAG-3’) (Riaz et al., 2011) using ecoPCR software (Ficetola et al., 2010). Set parameters allowed a 50-250 bp fragment and three mismatches between each primer and reference sequence. Primers were validated *in vitro* for UK fish by Hänfling et al. (2016) and for six UK amphibian species by Harper et al. (2018) (Fig. S2).

eDNA samples were amplified in triplicate with the aforementioned primers, where PCR positive controls (six per PCR plate; *n* = 114) were cichlid (*Rhamphochromis esox*) DNA (0.284 ng/µL) and PCR negative controls (six per PCR plate; *n* = 114) were sterile molecular grade water (Fisher Scientific UK Ltd, UK). PCR products were individually purified using E.Z.N.A^®^ Cycle Pure V-Spin Clean-Up Kits (Omega Bio-tek, GA, USA) following the manufacturer’s protocol. Purified product was pooled according to sample for the second PCR which bound Multiplex Identification tags to the purified products. PCR products were individually purified using magnetic bead clean-up and quantified with a Quant-IT™ PicoGreen™ dsDNA Assay (Invitrogen, UK). Samples were normalised, pooled, and libraries quantified using a Qubit™ dsDNA HS Assay (Invitrogen, UK). Libraries were sequenced on an Illumina® MiSeq using 2 × 300 bp V3 chemistry (Illumina, Inc, CA, USA) and raw sequence reads processed using metaBEAT (metaBarcoding and Environmental Analysis Tool) v0.97.7 (https://github.com/HullUni-bioinformatics/metaBEAT).

After quality filtering (phred score Q30) and trimming (length filter of 90-110 bp) using Trimmomatic v0.32 (Bolger, Lohse, & Usadel, 2014), merging (10 bp overlap minimum and 10% mismatch maximum) using FLASH v1.2.11 (Magoč & Salzberg, 2011), chimera detection using the uchime algorithm (Edgar et al., 2011) in vsearch v1.1 (Rognes et al., 2016), and clustering (97% identity with 3 sequences minimum per cluster) with vsearch v1.1 (Rognes et al., 2016), non-redundant query sequences were compared against the reference database using BLAST (Zhang et al., 2000). Putative taxonomic identity was assigned using a lowest common ancestor (LCA) approach based on the top 10% BLAST matches for any query matching with at least 98% identity to a reference sequence across more than 80% of its length. Unassigned sequences were subjected to a separate BLAST against the complete NCBI nucleotide (nt) database at 98% identity to determine the source via LCA as described above. The bioinformatic analysis was archived (https://doi.org/10.5281/zenodo.1188710) by Harper et al. (2018) for reproducibility.

### 2.3 Data analysis

#### 2.3.1 Dataset refinement

Analyses were performed in R v.3.4.3 (R Core Team, 2017). Data and R scripts have been deposited in a dedicated GitHub repository for this study, which has been permanently archived (https://doi.org/10.5281/zenodo.3365703). Assignments from different databases were merged, and spurious assignments (i.e. non-UK species, invertebrates and bacteria) removed from the dataset. The family Cichlidae was reassigned to *Rhamphochromis esox*. The green-winged teal (*Anas carolinenisis*) was reassigned to *Anas* (dabbling ducks) because this species is a rare migrant and available reference sequences were identical across our metabarcode to those for mallard (*Anas platyrhynchos*) and Eurasian teal (*Anas crecca*), which are widely distributed across the UK. Scottish wildcat (*Felis silvestris*) does not occur at the sampling localities (Kent, Lincolnshire and Cheshire) and was therefore reassigned to domestic cat (*Felis catus*). Wild boar (*Sus scrofa*) and grey wolf (*Canis lupus*) were reassigned to domestic pig (*Sus scrofa domesticus*) and domestic dog (*Canis lupus familiaris*) given the restricted distribution of *S. scrofa* and absence of *C. lupus* in the wild in the UK. The genus *Strix* was reassigned to tawny owl (*Strix aluco*) as it is the only UK representative of this genus. Where family and genera assignments containing a single UK representative had reads assigned to species, reads from all assignment levels were merged and manually assigned to that species.

Of the 114 PCR negative controls included, 50 produced no reads (Fig. S3). Reads generated for 64 of 114 PCR negative controls ranged from 0 to 49,227, and strength of each contaminant varied (mean = 0.021%, range = 0 – 100.0% of the total reads per PCR negative control). To minimise risk of false positives, we evaluated different sequence thresholds. These included the maximum sequence frequency of *R. esox* DNA in eDNA samples (100%), maximum sequence frequency of any DNA except *R. esox* in PCR positive controls (83.96%), and taxon-specific thresholds (maximum sequence frequency of each taxon in PCR positive controls). The different thresholds were applied to the eDNA samples and the results from each compared (Fig. S4). The taxon-specific thresholds (Table S3) retained the most biological information, thus these were selected for downstream analysis. Consequently, taxa were only classed as present at sites if their sequence frequency exceeded their threshold. After applying the taxon-specific thresholds, the PCR positive control (*R. esox*), human (*Homo sapiens*), and domestic species (Table S4) were removed from the dataset. Higher taxonomic assignments excluding the genus *Anas* were then removed, thus taxonomic assignments in the final dataset were predominantly of species resolution. The read count data were converted to a binary species matrix for downstream analyses using the package cooccur v1.3 (Griffith, Veech & Marsh, 2016) and Generalized Linear Mixed-effects Models (GLMMs), which were fit by maximum likelihood with the glmer function using the package lme4 v1.1-21 (Bates et al., 2015). County was included as a random effect in each GLMM to account for spatial dependencies between ponds (see Supporting Information: Appendix 1).

#### 2.3.2 Hypothesis generation: biotic factors potentially influencing T. cristatus detection

We first mined the eDNA metabarcoding data and investigated the influence of vertebrate group richness on *T. cristatus* detection. We used a binomial GLMM with the logit link function that included species richness of other amphibians, fish, waterfowl, terrestrial birds, and mammals as fixed effects (model 1, *N* = 532). We then performed a preliminary analysis using the package cooccur v1.3 (Griffith, Veech & Marsh, 2016) to identify species associations between *T. cristatus* and other vertebrates (*N* = 532). Hypotheses generated by the eDNA metabarcoding data relating to factors influencing *T. cristatus* detection are summarised in Table 1.

**Table 1.** Summary of hypothesis generation using eDNA metabarcoding data, specifically biotic factors that may influence *T. cristatus* detection probability. Direction of observed effects on eDNA-based *T. cristatus* detection identified by the GLMM assessing species richness in each vertebrate group (*N* = 532) and the preliminary cooccur analysis (*N* = 532) are given. Negative and positive effects are listed as – and + respectively. Test statistic is for LRT used and significant P-values (<0.05) are in bold.

| Variable | Analysis |  |  |  |  |  |  |  |
| --- | --- | --- | --- | --- | --- | --- | --- | --- |
|  | Cooccur |  |  |  | GLMM |  |  |  |
| | Effect | *P* | Adjusted *P*<br>(Benjamini-Hochberg) | DF | Effect size (SE) | $\chi^2$ | *P* | Adjusted *P*<br>(Benjamini-Hochberg) |
| **Amphibians** |  |  |  |  |  |  |  |  |
| Species richness |  |  |  | 1 | 0.526 (0.140) | 14.195 | **<0.001** | **<0.001** |
| *L. vulgaris* | + | **<0.001** | **<0.001** | 1 | 1.616 (0.232) |  |  |  |
| *B. bufo* | - | **0.010** | 0.489 | 1 | -1.793 (0.643) |  |  |  |
| **Fish** |  |  |  |  |  |  |  |  |
| Species richness |  |  |  | 1 | -0.163 (0.124) | 1.798 | 0.180 | 0.225 |
| *G. aculeatus* | - | **0.012** | 0.489 |  |  |  |  |  |
| *C. carpio* | - | **0.040** | 0.714 |  |  |  |  |  |
| *P. pungitius* | - | **0.049** | 0.714 |  |  |  |  |  |
| *C. carassius* |  |  |  |  |  |  |  |  |
| **Waterfowl** |  |  |  |  |  |  |  |  |
| Species richness |  |  |  | 1 | 0.491 (0.134) | 13.530 | **<0.001** | **<0.001** |
| *G. chloropus* | + | **0.001** | 0.012 |  |  |  |  |  |
| *F. atra* | + | **0.022** | 0.237 |  |  |  |  |  |
| **Terrestrial birds** |  |  |  |  |  |  |  |  |
| Species richness |  |  |  | 1 | -0.044 (0.302) | 0.021 | 0.884 | 0.884 |
|  | Cooccur |  |  |  | GLMM |  |  |  |
| | Effect | *P* | Adjusted *P*<br>(Benjamini-Hochberg) | DF | Effect size (SE) | $\chi^2$ | *P* | Adjusted *P*<br>(Benjamini-Hochberg) |
| *P. colchicus* | - | **0.050** | 0.714 |  |  |  |  |  |
| **Terrestrial mammals** |  |  |  |  |  |  |  |  |
| Species richness |  |  |  | 1 | -0.515 (0.188) | 8.501 | **0.004** | **0.006** |
| *S. carolinensis* | - | **0.020** | 0.685 |  |  |  |  |  |

#### 2.3.3 Hypothesis testing: biotic and abiotic factors known to influence T. cristatus detection

Identified associations from the cooccur analysis in conjunction with the existing *T. cristatus* literature informed candidate biotic variables to be modelled against *T. cristatus* detection (model 2, *n* = 504). The existing *T. cristatus* literature informed candidate abiotic variables to be modelled against *T. cristatus* detection (model 3, *n* = 504). Hypotheses tested using the eDNA metabarcoding data and associated environmental metadata to examine biotic and abiotic factors influencing *T. cristatus* detection are summarised in Tables 2 and S5. Selection of a suitable set of explanatory variables and modelling framework is fully described in Supporting Information: Appendix 1. Briefly, candidate biotic and abiotic explanatory variables were assessed for: collinearity using a Spearman’s rank pairwise correlation matrix and examination of variance inflation factors (Zuur et al., 2009), relative importance using a classification tree within the package rpart v4.1-13 (Therneau, Atkinson & Ripley, 2014), and non-linearity using Generalized Additive Models (GAMs) (Zuur et al., 2009). We constructed separate binomial GLMMs with the logit link function for biotic (model 2) and abiotic (model 3) explanatory variables that passed these assessments.

#### 2.3.4 Hypothesis testing: T. cristatus HSI and umbrella status

We modelled HSI score (fixed effect) separately to prevent HSI score masking variation caused by the individual biotic and abiotic variables it encompasses (model 4). Using a Poisson GLMM, we tested the umbrella species status of *T. cristatus* by modelling vertebrate species richness against *T. cristatus* detection and the *T. cristatus* HSI score (model 5).

#### 2.3.5 Model selection and validation

For each GLMM, we employed an information-theoretic approach using Akaike Information Criterion (AIC) to determine the most parsimonious approximating model to make predictions (Akaike, 1973). Biotic and abiotic models considered respectively (Table S5) were nested thus the best models were chosen using stepwise backward deletion of terms based on Likelihood Ratio Tests (LRTs). Models 1, 4 and 5 were compared to null GLMMs. The final models resulting from stepwise selection were as follows:

Model 1: *T. cristatus* ∼ (1|County) + amphibian richness + fish richness + waterfowl richness+ terrestrial bird richness + mammal richness

Model 2: *T. cristatus* ∼ (1|County) + *L. vulgaris* + *B. bufo* + *C. carpio* + *G. aculeatus* + *Gallinula chloropus* + Waterfowl

Model 3: *T. cristatus* ∼ (1|County) + inflow + pond area + shading

Model 4: *T. cristatus* ∼ (1|County) + HSI score

Model 5: Vertebrate species richness ∼ (1|County) + *T. cristatus* + HSI score

Final models were tested for overdispersion using a custom function testing overdispersion of the Pearson residuals. Model fit was assessed using the Hosmer and Lemeshow Goodness of Fit Test within the package ResourceSelection v0.3-2 (Lele, Keim & Solymos, 2016), quantile-quantile plots, and partial residual plots (Zuur et al., 2009). Conditional *R*^2^ was calculated using the r.squaredGLMM function in the package MuMIn v1.42.1 (Bartoń, 2018). The Benjamini-Hochberg correction was calculated for explanatory variable p-values from the cooccur and GLMM analyses using the p.adjust function in the package stats v3.4.3, and this alpha applied to the results to account for potential Type I error. The original p-values and the adjusted Benjamini-Hochberg p-values are reported for comparison in Tables 1 and 2. Model predictions were obtained using the predictSE function in the package AICcmodavg v2.1-1 (Mazerolle, 2016) and upper and lower 95% CIs were calculated from the standard error of the predictions. Results were plotted using the package ggplot2 v3.1.0 (Wickham, 2016).

**Table 2.**
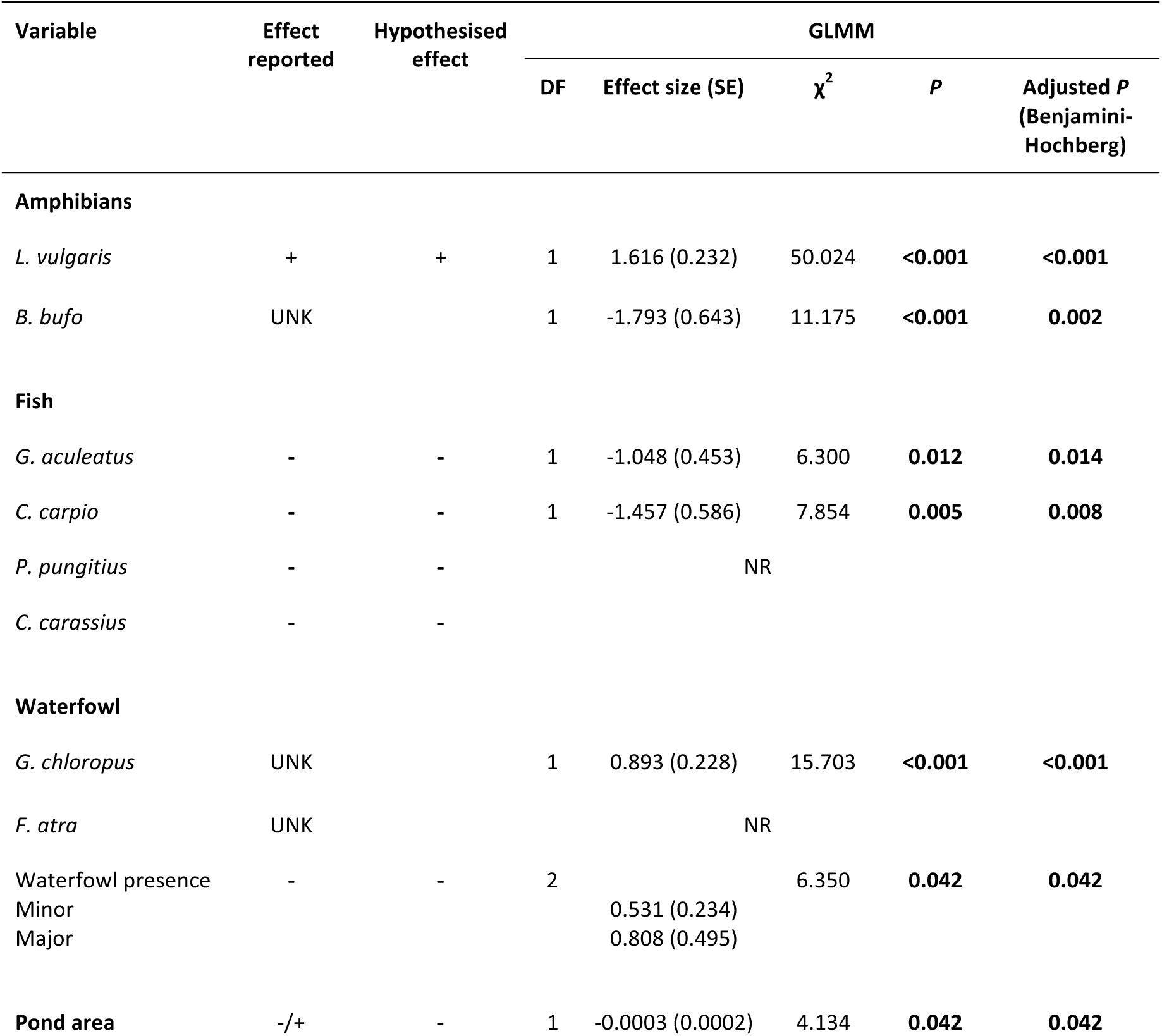
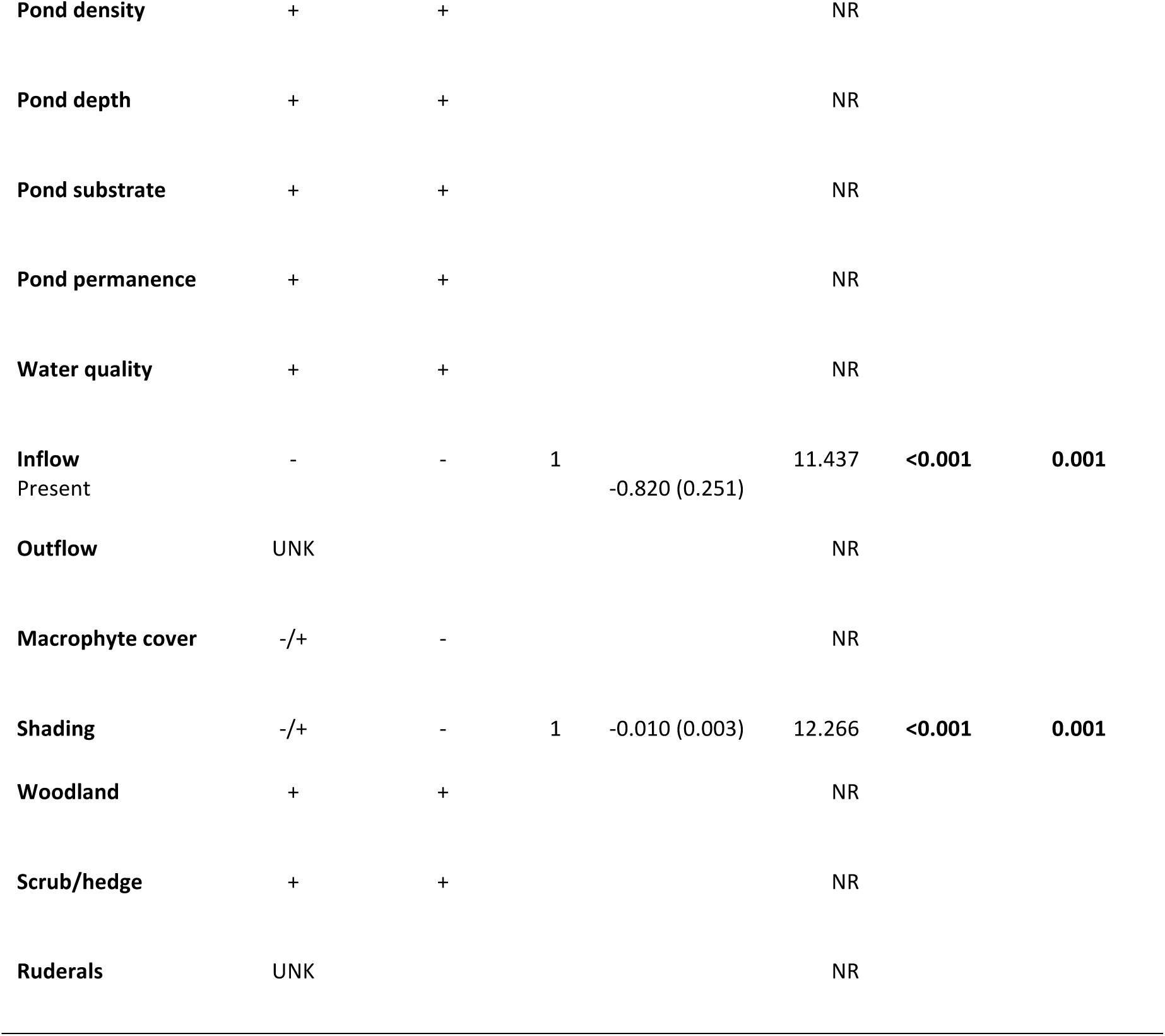
Summary of hypothesis testing using eDNA metabarcoding data and associated environmental metadata, specifically biotic and abiotic factors known to influence *T. cristatus* detection probability. Reported effects on *T. cristatus* detection in the literature and hypothesised effects on eDNA-based *T. cristatus* detection are given for each variable. Any variables not explicitly reported in the literature that were identified by the preliminary cooccur analysis are listed as UNK. Direction of observed effects on eDNA-based *T. cristatus* detection identified by each analysis (biotic GLMM, *n* = 504; abiotic GLMM, *n* = 504) are given. Negative and positive effects are listed as – and + respectively. For categorical variables with more than one level, effect size and standard error (SE) are only given for levels reported in the model summary. Test statistic is for LRT used and significant P-values (<0.05) are in bold. Variables included for model selection but not retained in the final models are listed as NR.

## 3. Results

### 3.1 eDNA metabarcoding

Harper et al. (2018) processed a total of 532 eDNA samples and 228 PCR controls across two sequencing runs. The runs generated raw sequence read counts of 36,236,862 and 32,900,914 respectively. After quality filtering, trimming, and merging of paired-end reads, 26,294,906 and 26,451,564 sequences remained. Following removal of chimeras and read dereplication via clustering, the libraries contained 14,141,237 and 14,081,939 sequences (average read counts of 36,826 and 36,671 per sample respectively), of which 13,126,148 and 13,113,143 sequences were taxonomically assigned. The final dataset used here (assignments corrected, thresholds applied, and assignments removed) contained 53 vertebrate species (Table S6), including six amphibians, 14 fish, 17 birds, and 16 mammals (Fig. 1).

**Figure 1.**
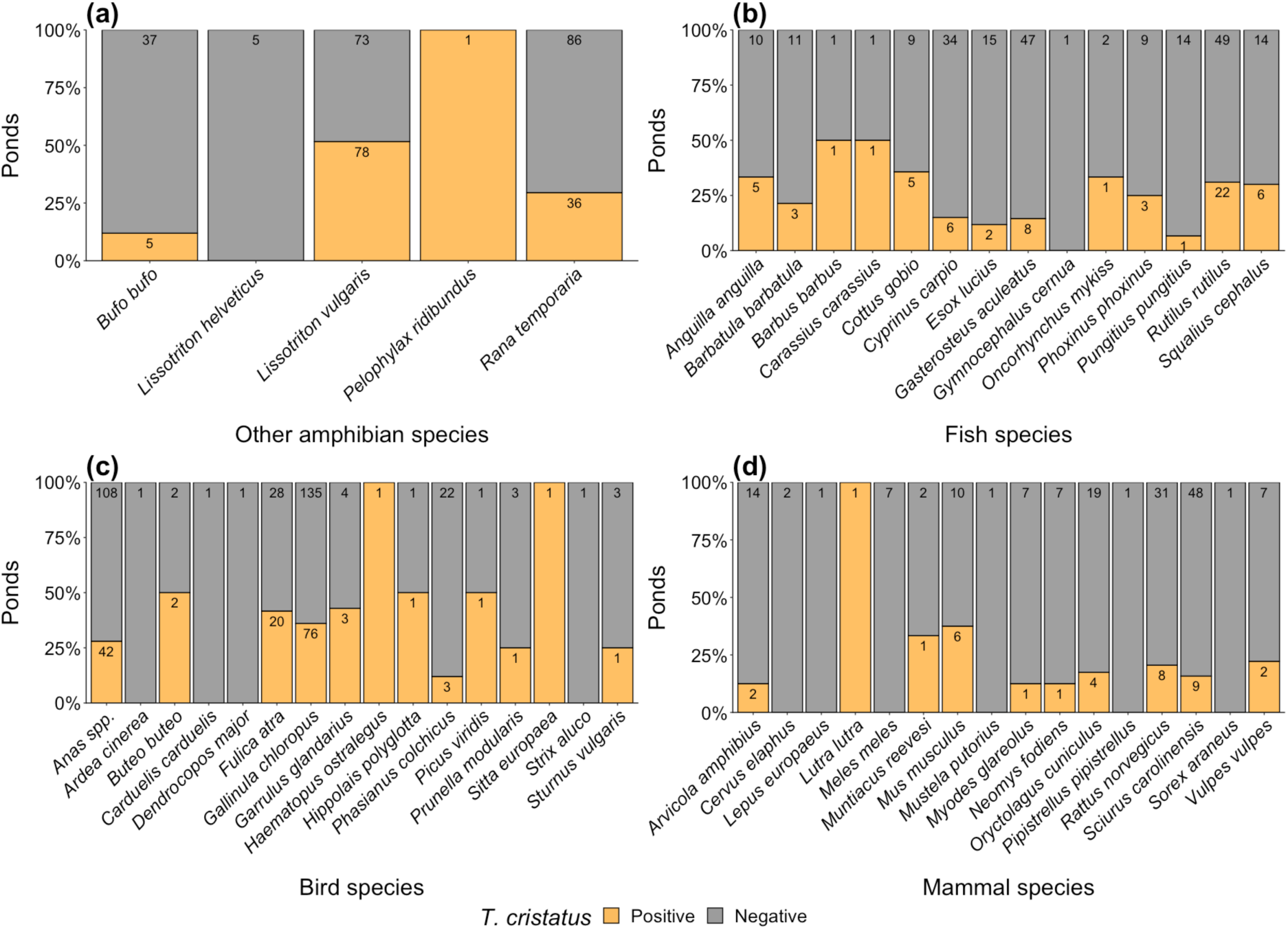
eDNA metabarcoding detection of *T. cristatus* in relation to other vertebrate species (*N* = 532 ponds): **(a)** other amphibians, **(b)** fish, **(c)** birds, and **(d)** mammals. Numbers on each bar are the number of ponds with (orange) and without (grey) *T. cristatus* in which a vertebrate species was detected.

### 3.2 Pondscape biodiversity

All native amphibians were detected as well as the non-native marsh frog (*Pelophylax ridibundus*). *T. cristatus* (*n* = 148), *L. vulgaris* (*n* = 151) and *R. temporaria* (*n* = 122) were widespread, but *B. bufo* (*n* = 42), palmate newt (*Lissotriton helveticus* [*n* = 5]) and *P. ridibundus* were uncommon (*n* = 1). The threatened European eel (*Anguilla anguilla* [*n* = 15]), European bullhead (*Cottus gobio* [*n* = 14]), and *C. carassius* (*n* = 2) were detected alongside native fishes, such as pike (*Esox Lucius* [n = 17]) and roach (*Rutilus rutilus* [*n* = 71]), but also introduced species, including *C. carpio* (*n* = 40), ruffe (*Gymnocephalus cernua* [*n* = 1]), and rainbow trout (*Oncorhynchus mykiss* [*n* = 3]). Some identified waterfowl were ubiquitous, such as common moorhen (*Gallinula chloropus* [*n* = 211]), whereas others were less common, e.g. grey heron (*Ardea cinerea* [*n* = 1]) and Eurasian oystercatcher (*Haematopus ostralegus* [*n* = 1]). Terrestrial fauna were often detected in fewer than five ponds (Figs. 1c, d). Buzzard (*Buteo buteo* [*n* = 4]), Eurasian jay (*Garrulus glandarius* [*n* = 7]), dunnock (*Prunella modularis* [*n* = 4]), and starling (*Sturnus vulgaris* [*n* = 4]) were the most frequently detected terrestrial birds. Introduced mammals (Mathews et al., 2018), such as grey squirrel (*Sciurus carolinensis* [*n* = 57]) and Reeve’s muntjac (*Muntiacus reevesi* [*n* = 3]), outweighed native mammals. Nonetheless, we detected several mammals with Biodiversity Actions Plans and/or of conservation concern (Mathews et al., 2018), including otter (*Lutra lutra* [*n* = 1]), water vole (*Arvicola amphibius* [*n* = 16]), European polecat (*Mustela putorius* [*n* = 1]), brown hare (*Lepus europaeus* [*n* = 1]) and water shrew (*Neomys fodiens* [*n* = 8]). Notably, the invasive American mink (*Neovison vison*) was not detected despite widespread UK distribution (Mathews et al., 2018). All species and their detection frequencies are listed in Table S6.

### 3.3 Hypothesis generation: biotic factors potentially influencing *T. cristatus* detection

*T. cristatus* detection was positively associated with amphibian and waterfowl species richness, yet negatively associated with mammal species richness (Table 1). *T. cristatus* detection was less probable as fish and terrestrial bird species richness increased, but these trends were not significant (Table 1, Fig. 2; GLMM: overdispersion θ = 0.995, *χ*^2^_525_ = 522.276, *P* = 0.525; fit *χ*^2^_8_ = 1.568, *P* = 0.992, *R*^2^ = 12.74%). With no correction for Type I error, *T. cristatus* had significant positive associations with three species (Table 2, Fig. S5), including *L. vulgaris*, common coot (*Fulica atra*) and *G. chloropus*, and significant negative associations with six species (Table 2, Fig. S5), including *B. bufo, C. carpio, G. aculeatus, P. pungitius*, common pheasant (*Phasianus colchicus*) and *S. carolinensis*. However, only *T. cristatus* associations with *L. vulgaris* and *G. chloropus* were retained after applying the Benjamini-Hochberg correction (Table 2).

**Figure 2.**
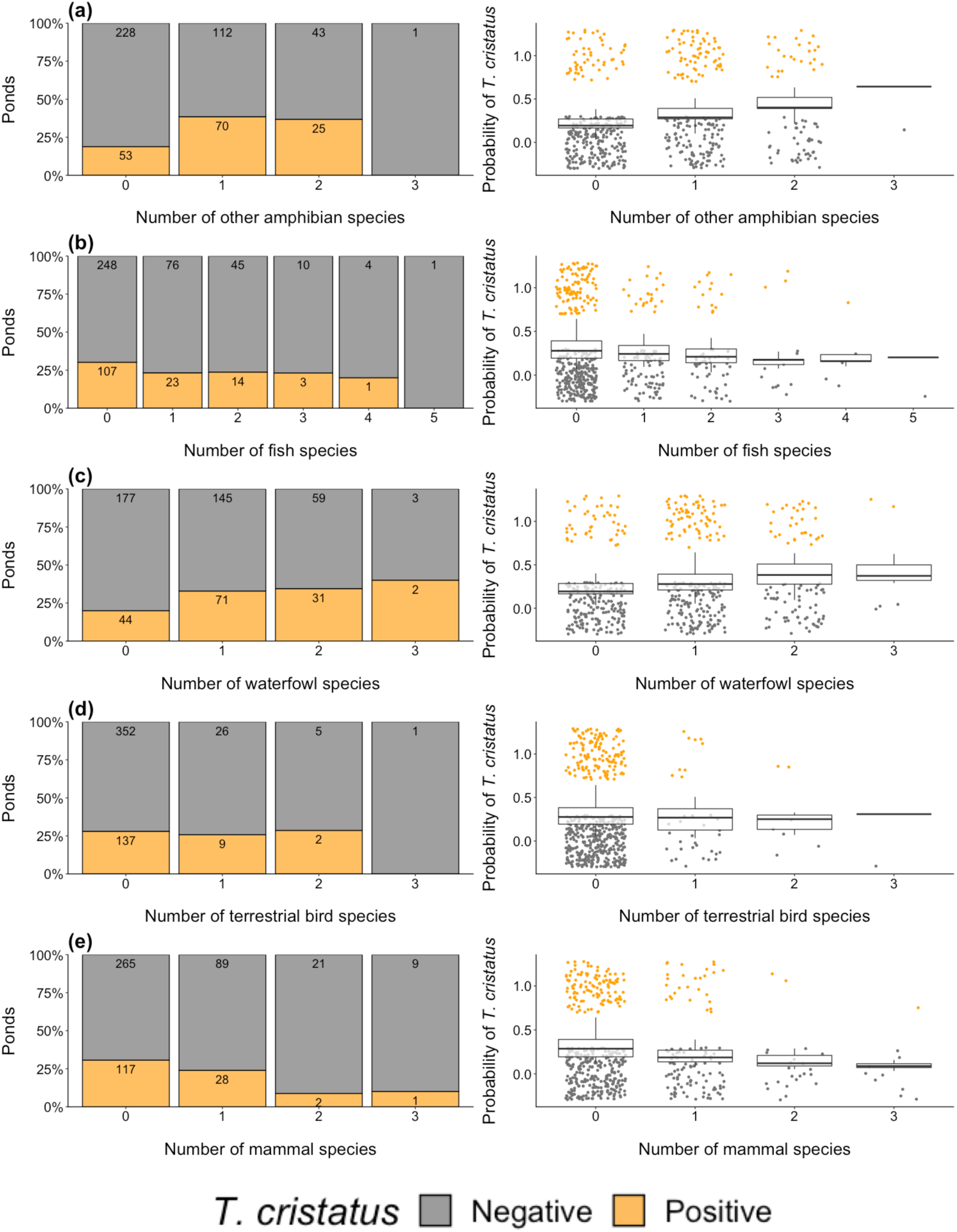
*T. cristatus* detection (orange) and non-detection (grey) in relation to species richness of different vertebrate groups (*N* = 532 ponds): **(a)** other amphibians, **(b)** fish, **(c)** waterfowl, **(d)** terrestrial birds, and **(e)** mammals. Observed proportion of ponds with and without *T. cristatus* (left) is plotted alongside predicted probability of *T. cristatus* detection (right). Numbers on barplots of observed detection are the number of ponds for each category. In plots showing predicted *T. cristatus* detection, the observed data is shown as points (jittered around 0 and 1 to clarify variation in point density) and boxes are the model predictions.

*T. cristatus* detection was more likely in ponds with more amphibian species (Table 1, Fig. 2a). *T. cristatus* was detected in 51.66% of ponds (*n* = 151) identified as containing *L. vulgaris*, but in only 11.91% of ponds (*n* = 42) where *B. bufo* was detected (Fig. 1a). Probability of *T. cristatus* detection was lower in ponds with more fish species (Table 1), and *T. cristatus* was not detected in ponds with more than four fish species (Fig. 2b). *T. cristatus* was only detected in 15.00% (*n* = 40), 14.55% (*n* = 55) and 6.67% (*n* = 15) of ponds identified as containing *C. carpio, G. aculeatus* and *P. pungitius* respectively (Fig. 1b). In contrast, *T. cristatus* detection was more likely in ponds where more waterfowl species were detected (Table 1, Fig. 2c). *T. cristatus* was detected in 41.67% (*n* = 48) and 36.02% (*n* = 211) of ponds identified as containing *F. atra* and *G. chloropus* respectively (Fig. 1c). *T. cristatus* detection was negatively correlated with higher terrestrial bird species richness, but not significantly so (Table 1, Fig. 2d). However, *T. cristatus* was only detected in 12.00% (*n* = 25) ponds where *P. colchicus* was detected (Fig. 1c). *T. cristatus* detection was less likely in ponds where more mammal species were detected (Table 1, Fig. 2e). Specifically, *T. cristatus* was only detected in 15.79% (*n* = 57) of ponds where *S. carolinensis* was detected (Fig. 1d).

### 3.4 Hypothesis testing: biotic and abiotic factors known to influence *T. cristatus* detection

Biotic variables put forward for model selection were eDNA-based detection/non-detection of *L. vulgaris, B. bufo, C. carpio, G. aculeatus, P. pungitius, G. chloropus*, and *F. atra* as well as presence/absence of other amphibians, fish, and waterfowl as recorded during HSI assessment by environmental consultants (Table S5). Only detection/non-detection of *L. vulgaris, B. bufo, C. carpio, G. aculeatus*, and *G. chloropus* as well as waterfowl presence were retained by model selection as explanatory variables for the biotic GLMM of *T. cristatus* detection (Table 2, Figs. 3a-f; GLMM: overdispersion θ = 1.013, *χ*^2^_495_ = 495.000, *P* = 0.411; fit *χ*^2^_8_ = 4.437, *P* = 0.8157, *R*^2^ = 27.77%). Abiotic variables put forward for model selection were max. depth, pond density, presence of inflow, pond area, pond substrate, presence of outflow, percentage of macrophyte cover, water quality, pond permanence, percentage of shading, ruderals, scrub/hedge, and woodland (Table S5). Only presence of inflow, pond area, and percentage of shading were retained by model selection as explanatory variables for the abiotic GLMM explaining *T. cristatus* detection (Table 2, Figs. 3g-i; GLMM: overdispersion θ = 1.000, *χ*^2^_499_ = 499.126, *P* = 0.490; fit *χ*^2^_8_ = 4.898, *P* = 0.768, *R*^2^ = 10.00%). Results are summarised and relationships compared to those previously reported for *T. cristatus* in Table 2. Application of the Benjamini-Hochberg correction for Type I error did not alter variable significance.

**Figure 3.**
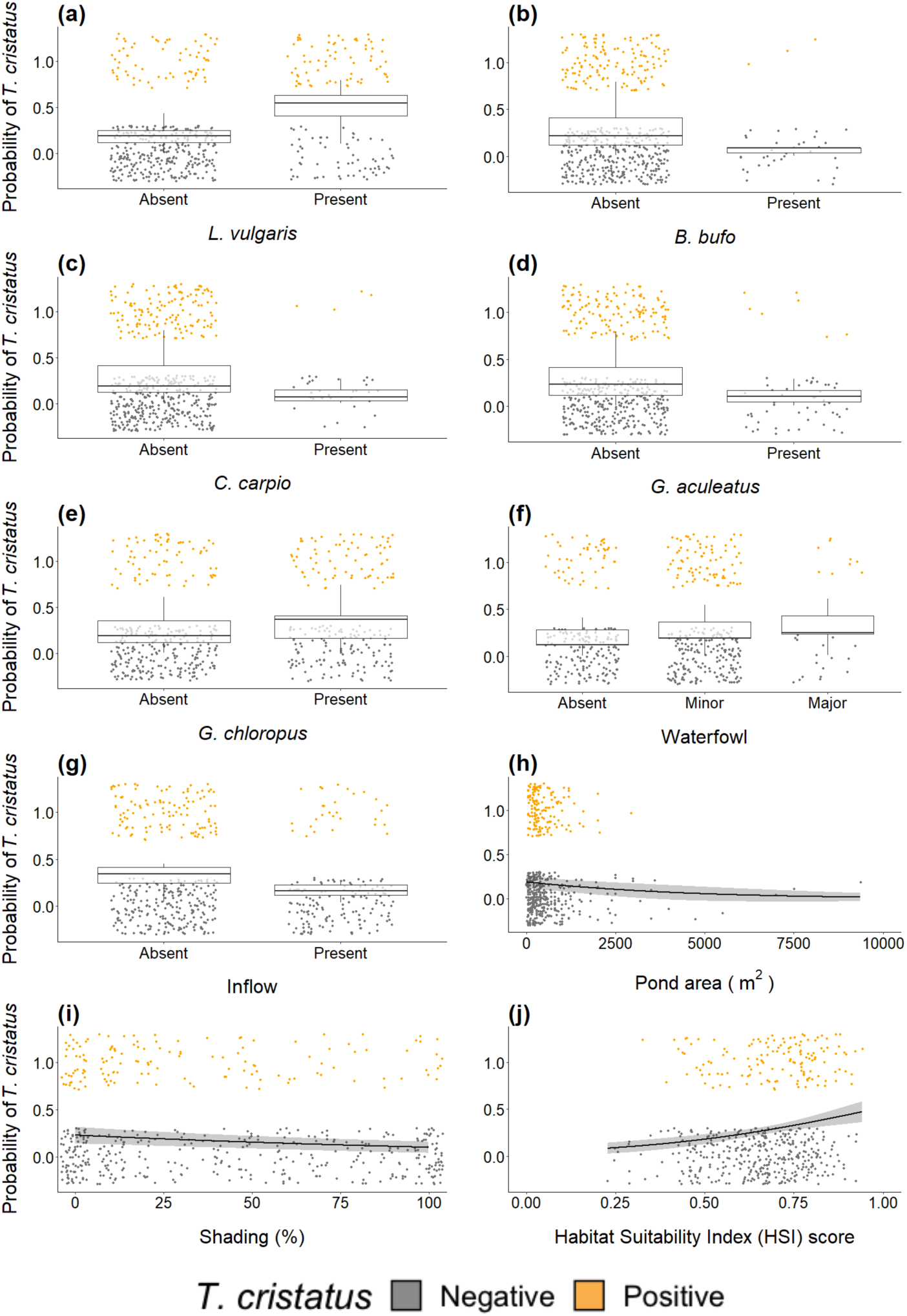
Biotic and abiotic factors influencing *T. cristatus* detection (*n* = 504 ponds): **(a)** *L. vulgaris* detection, **(b)** *B. bufo* detection, **(c)** *C. carpio* detection, **(d)** *G. aculeatus* detection, **(e)** *G. chloropus* detection, **(f)** extent of waterfowl presence, **(g)** presence of inflow, **(h)** pond area, **(i)** percentage of shading, and **(j)** HSI score. The 95% CIs, as calculated using the predicted *T. cristatus* probability values and standard error for these predictions, are given for each relationship. The observed *T. cristatus* detection (orange) and non-detection (grey) data are displayed as points (jittered around 0 and 1 to clarify variation in point density) against the predicted relationships (boxes/lines).

### 3.5 Hypothesis testing: *T. cristatus* HSI and umbrella status

HSI score positively correlated with *T. cristatus* detection probability (GLMM: overdispersion θ = 0.995, *χ*^2^_501_ = 498.372, *P* = 0.525; fit *χ*^2^_8_ = 5.395, *P* = 0.715, *R*^2^ = 6.66%), where *T. cristatus* detection was more likely in ponds with a higher HSI score (Fig. 3j; estimate ± standard error = 3.094 ± 0.803, *χ*^2^_1_ = 16.020, *P* < 0.001). Vertebrate species richness was positively associated with *T. cristatus* detection (GLMM: overdispersion θ = 1.112, *χ*^2^_500_ = 555.800, *P* = 0.042; fit *χ*^2^_8_ = −11.91, *P* = 1.000, *R*^2^ = 9.14%), with more species detected in ponds identified as containing *T. cristatus* (Fig. 4a; estimate ± standard error = 0.408 ± 0.059, *χ*^2^_1_ = 46.265, *P* < 0.001, adjusted *P* [Benjamini-Hochberg] < 0.001). However, vertebrate species richness did not significantly increase with the *T. cristatus* HSI score (Fig. 4b; estimate ± standard error = 0.228 ± 0.206, *χ*^2^_1_ = 1.227, *P* = 0.268, adjusted *P* [Benjamini-Hochberg] = 0.268).

**Figure 4.**
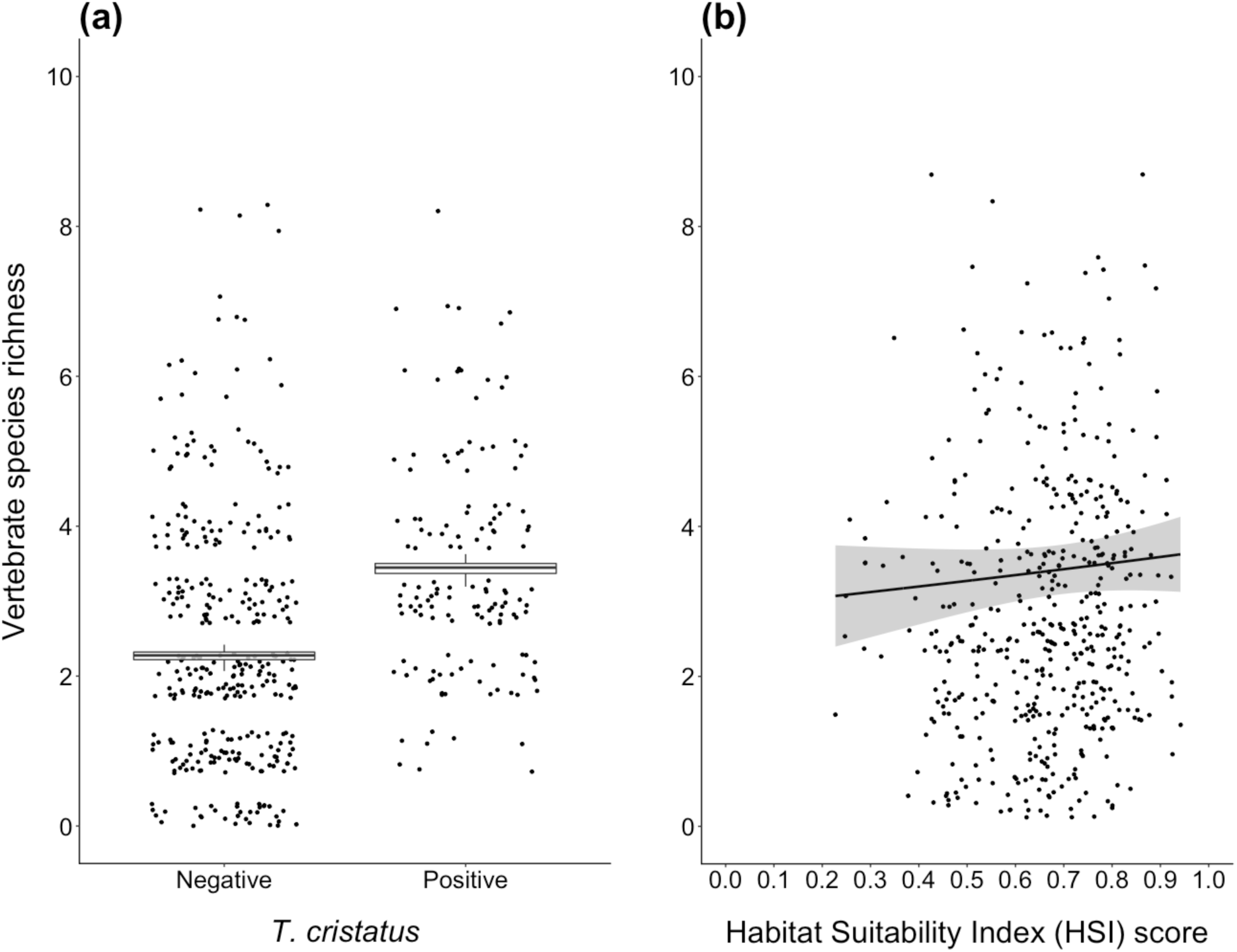
Vertebrate species richness in ponds (*n* = 504) in relation to: **(a)** *T. cristatus* detection and **(b)** the *T. cristatus* HSI score. The 95% CIs, as calculated using the predicted species richness values and standard error for these predictions, are given for each relationship. The observed data are displayed as points (jittered to clarify variation in point density) against the predicted relationships (boxes/lines).

## 4. Discussion

We have validated eDNA metabarcoding for ecological hypothesis generation and testing using the community data produced by this tool in combination with environmental metadata for ponds. We identified and tested biotic and abiotic factors influencing *T. cristatus* detection, whether the HSI can be applied to *T. cristatus* eDNA survey, and whether *T. cristatus* is truly an umbrella species for pond conservation. *T. cristatus* detection was more likely in ponds where *L. vulgaris* and *G. chloropus* were detected, and ponds where *B. bufo, C. carpio*, and *G. aculeatus* were not detected. *T. cristatus* detection was also more likely in ponds where waterfowl presence was recorded during HSI assessment. Ponds identified as containing *T. cristatus* were typically small, absent of inflow, and not excessively shaded. The *T. cristatus* HSI correlated with *T. cristatus* detection, but not vertebrate species richness. Nonetheless, more vertebrates were detected in ponds where *T. cristatus* was detected thus presence of this amphibian may indicate good quality habitat for other vertebrates. Our findings demonstrate the power of eDNA metabarcoding to enhance freshwater monitoring and research by providing biodiversity data *en masse* at low cost.

### 4.1 Pondscape biodiversity

eDNA metabarcoding detected six amphibian, 14 fish, 17 bird, and 16 mammal species across 532 UK ponds. This diverse species inventory emphasises the importance of ponds as habitat for aquatic taxa, but also as stepping stones for semi-aquatic and terrestrial taxa (De Meester et al., 2005; Hill et al., 2018) through provision of drinking, foraging, dispersive, and reproductive opportunities (Biggs et al., 2016; Klymus et al., 2017). Some species detections may be the result of eDNA transport from water bodies in the surrounding area (Hänfling et al., 2016) to ponds via inflow. However, this signifies the capacity of ponds to provide natural samples of freshwater and terrestrial biodiversity in the wider catchment (Deiner et al., 2017).

### 4.2 Biotic factors influencing *T. cristatus* detection

*T. cristatus* detection was more likely in ponds with higher amphibian species richness – particularly ponds identified as containing *L. vulgaris* but not *B. bufo. T. cristatus* and *L. vulgaris* have similar habitat requirements and tend to breed in the same ponds (Skei et al., 2006; Rannap et al., 2009a; Denoël et al., 2013; Cayuela et al., 2018), with >60% overlap reported (Rannap & Briggs, 2006). However, *L. vulgaris* can inhabit a broader range of habitat (Rannap & Briggs, 2006; Skei et al., 2006) than *T. cristatus*, which depends on larger, deeper ponds with abundant macrophytes and no fish located in open, semi-rural landscapes (Denoël et al., 2013). *B. bufo* can inhabit fish-containing ponds (Manenti & Pennati, 2016) and *T. cristatus* may predate *B. bufo* eggs and larvae (Langton et al., 2001). This may explain the negative association between *B. bufo* and *T. cristatus* as opposed to the positively associated *T. cristatus* and *L. vulgaris*.

*T. cristatus* detection marginally decreased with higher fish species richness, and *T. cristatus* was negatively associated with *C. carpio, G. aculeatus*, and *P. pungitius*. These fishes are common in and typical of ponds. All *T. cristatus* life stages may be predated by fishes (Langton et al., 2001) and negative effects of fish presence-absence on *T. cristatus* occupancy, distribution, and abundance are repeatedly reported (Joly et al., 2001; Rannap & Briggs, 2006; Skei et al., 2006; Denoël & Ficetola, 2008; Rannap et al., 2009a, b; Hartel et al., 2010; Denoël et al., 2013). *G. aculeatus* predates *T. cristatus* eggs and larvae (McLee & Scaife, 1992; Jarvis, 2010), and has non-consumptive effects on *T. cristatus* embryos (Jarvis, 2010). *T. cristatus* larvae were also found to alter their behaviour when exposed to predatory *G. aculeatus* but not non-predatory *C. carassius* (Jarvis, 2012), another fish characteristic of ponds.

In our study, we detected *T. cristatus* in 50% of ponds where *C. carassius* was detected, but <20% of ponds where large and/or predatory fishes were detected, e.g. *C. carpio, G. aculeatus, E. lucius*. Although we observed fewer detections from ponds for *C. carassius* than *C. carpio, G. aculeatus* or *E. lucius*, previous research also indicates large and/or predatory fish are more detrimental to *T. cristatus* occurrence (Skei et al., 2006; Hartel et al., 2010; Chan, 2011). *C. carassius* does not hinder *T. cristatus* oviposition, larval behaviour, or recruitment success (Chan, 2011; Jarvis, 2012), or pond invertebrate and macrophyte diversity (Stefanoudis et al., 2017). In contrast, *C. carpio* foraging reduces invertebrate density and macrophyte cover (Maceda-Veiga, López & Green, 2017), which lowers *T. cristatus* reproductive and foraging success and heightens predator exposure (Rannap & Briggs, 2006; Gustafson et al., 2006; Chan, 2011). *C. carassius* and *C. carpio* are both included among fish species assumed to negatively impact *T. cristatus* and whose presence-absence is assessed for the *T. cristatus* HSI (ARG-UK, 2010). However, given that *C. carassius* does not directly predate *T. cristatus* or indirectly alter its behaviour, reproductive success, or habitat, we advocate a systematic re-evaluation of problematic fish species for *T. cristatus* conservation.

*T. cristatus* detection was positively associated with waterfowl species richness, namely *F. atra* and *G. chloropus* detection. These waterfowl species share macrophytes and macroinvertebrates as resources with amphibians, feeding on both directly (Perrow et al., 1997; Paillisson & Marion, 2001; Wallau et al., 2010). *F. atra* and *G. chloropus* crop emergent macrophytes to search for invertebrate prey (Paillisson & Marion, 2001; Wallau et al., 2010), which may indirectly benefit *T. cristatus* foraging. Although *Fulica* spp. can also pull up submerged vegetation and damage vegetation banks (Lauridsen, Jeppesen & Andersen, 1993), diet is macrophyte-dominated in late summer and autumn (Perrow et al., 1997) and unlikely to impact *T. cristatus* breeding in spring (Langton et al., 2001). The positive association identified here between *T. cristatus* and these waterfowl most likely reflects a shared preference for macrophyte-rich ponds.

*T. cristatus* detection was less frequent in ponds with higher mammal species richness. Our preliminary cooccur analysis indicated *T. cristatus* had negative associations with *P. colchicus* and *S. carolinensis*, but not when a correction for Type I error was applied. *T. cristatus* associations with *F. atra, B. bufo, C. carpio, G. aculeatus*, and *P. pungitius* were also non-significant when this correction was applied. These associations require further investigation to determine whether they are real, and if so, whether they are direct and reflect competition, predation or shared resources as discussed above, or indirect and reflect land-use or effects of additional species.

### 4.3 Abiotic factors influencing *T. cristatus* detection

*T. cristatus* detection was less likely in large ponds with inflow present and a greater percentage of shading. Effects of pond area may depend on the size range of ponds studied. Although our results indicate *T. cristatus* prefers smaller ponds, pond area does not always influence occupancy (Maletzky, Kyek & Goldschmid, 2007; Denoël & Ficetola, 2008; Gustafson et al., 2011) and was deemed a poor predictor of reproductive success (Vuorio et al., 2013). *T. cristatus* has been found to utilise small and large ponds (Rannap & Briggs, 2006; Skei et al., 2006); however, very small ponds (<124 m^2^) may be unable to support all life stages, and larger ponds may contain fish and experience eutrophication due to agricultural or polluted run-off (Rannap & Briggs, 2006). Inflow to ponds may exacerbate these problems by facilitating entry of agricultural or polluted run-off and connections to streams and rivers containing large, predatory fish (Freshwater Habitats Trust, 2015). Our results corroborate existing research where viable *T. cristatus* populations were unlikely in ponds that were shaded (Vuorio et al., 2013) or had dense macrophyte cover (Rannap & Briggs, 2006; Skei et al., 2006; Hartel et al., 2010).

In our study, most environmental metadata available were qualitative, preventing detailed analyses on pond properties and terrestrial habitat in relation to *T. cristatus* detection. Better understanding of *T. cristatus* detection in relation to species interactions and habitat quality could be achieved with quantitative data on pond properties (e.g. water chemistry), terrestrial habitat (e.g. type, density, distance to ponds), and aquatic and terrestrial habitat usage by different vertebrate species. Furthermore, given the metapopulation dynamics of *T. cristatus*, future research should investigate spatial drivers (e.g. pond density and other indices of connectivity, land cover, climate variables, roads, rivers, elevation) of *T. cristatus* detection using innovative modelling approaches, such as individual-based models (Messager & Olden, 2018). However, acquiring this data to perform these models is a phenomenal task for large numbers of ponds across a vast landscape (Denoël & Ficetola, 2008).

### 4.4 *T. cristatus* HSI and umbrella status

We found the HSI can predict eDNA-based *T. cristatus* detection at the UK pondscape. This contradicts conventional studies which deemed the index inappropriate for predicting *T. cristatus* occupancy or survival probabilities (Unglaub et al., 2015). We detected more vertebrates in ponds identified as containing *T. cristatus*, which may support its status as an umbrella species for pond biodiversity and conservation (Gustafson et al., 2006). We also observed a non-significant increase in vertebrate species richness with increasing *T. cristatus* HSI score. An adapted HSI, designed to predict species richness, could help select areas for management and enhancement of aquatic and terrestrial biodiversity. Until then, presence of *T. cristatus* and its HSI may confer protection to broader biodiversity by identifying optimal habitat for pond creation and restoration to encourage populations of this threatened amphibian. The HSI is not without issue due to qualitative data used for score calculation and subjective estimation of indices (Oldham et al., 2000; O’Brien et al., 2017). For future application of this index in *T. cristatus* eDNA survey, we recommend metabarcoding to quantify some qualitatively assessed indices (e.g. water quality via macroinvertebrate diversity, fish and waterfowl presence) alongside *T. cristatus* detection. Provided rigorous spatial and temporal sampling are undertaken, eDNA metabarcoding can also generate site occupancy data to estimate relative species abundance (Valentini et al., 2016; Hänfling et al., 2016; Lawson Handley et al., 2019; Li et al., 2019a).

### 4.5 Limitations of repurposed eDNA samples for metabarcoding applications

This study was based on samples that were repurposed from targeted eDNA surveys for *T. cristatus*. eDNA sampling, capture, and extraction was conducted in accordance with Biggs et al. (2015), whose methods were chosen based on the eDNA literature at that time. Six 15 ml water samples were taken from a homogenised sample (600 ml). Ethanol precipitation was used for eDNA capture, followed by DNA extraction on the combined lysate from the six subsamples. Consequently, ponds were represented by a single eDNA sample, for which 12 qPCR replicates were performed (Biggs et al., 2015). However, eDNA metabarcoding may require an entirely different workflow to enhance species detection (Harper et al., 2018; Harper et al., 2019).

Independent biological replicates as opposed to pseudoreplicates from a single water sample are key to account for spatial heterogeneity of eDNA from different species (Hänfling et al., 2016; Bálint et al., 2018; Lawson Handley et al., 2019; Li et al., 2019b). Larger volumes should be filtered instead of ethanol precipitation on small volumes to maximise eDNA capture (Harper et al., 2019). Samples should be extracted individually to maintain biological replication followed by independent technical replicates for PCR and sequencing. These levels of replication are required for hierarchical occupancy modelling to identify false positives and species detection probabilities as well as minimise false negatives (Ficetola et al., 2015; Bálint et al., 2018; Dorazio & Erickson, 2018).

We performed three PCR replicates for each sample that were pooled prior to sequencing. We included a large number of PCR controls and applied sequence thresholds to control for any false positives in our dataset arising from potential laboratory contamination. Pooling of technical replicates allowed us to screen vertebrate communities from over 500 ponds, but prevented occupancy modelling to verify species detection. Samples collected for targeted eDNA analysis can vary widely in terms of sampling strategy, capture, extraction and storage protocols. Ultimately, these protocols may limit the data generated by eDNA metabarcoding and the analyses that can be performed. This is an important consideration for any future research where eDNA samples are repurposed to address different ecological questions.

### 4.6 Prospects of eDNA metabarcoding for freshwater conservation, management, and research

We have demonstrated the effectiveness of eDNA metabarcoding for landscape-scale biodiversity monitoring as well as ecological hypothesis generation and testing. We combined metabarcoding with environmental metadata to identify new and revisit old hypotheses relating to biotic and abiotic factors that influence a threatened amphibian at the UK pondscape. Our findings will guide *T. cristatus* conservation in the face of increasing land-use and habitat fragmentation – a poignant issue as protective legislation for this species in the UK is changing. Whilst conservation of threatened species and their habitat should be a priority, the bigger picture should not be ignored. eDNA metabarcoding could enhance our understanding of freshwater networks, particularly pondscapes, to enable more effective monitoring, protection, and management of aquatic and terrestrial biodiversity. We are only now beginning to realise and explore these opportunities.

## Acknowledgements

This work was funded by the University of Hull. We would like to thank Jennifer Hodgetts (Fera Science Ltd) for assisting with sample collection, and Jianlong Li (University of Hull) for sequencing primer design and advice on laboratory protocols. Tissue samples for primer validation and Sanger sequencing were provided by Andrew Buxton and Richard Griffiths (DICE, University of Kent) under licence from Natural England, and Barbara Mabel and Elizabeth Kilbride (University of Glasgow).

## Data Availability

The taxonomically assigned sequence reads used in this study were generated by Harper et al. (2018). The raw sequence reads were archived on the NCBI Sequence Read Archive (Bioproject: PRJNA417951; SRA accessions: SRR6285413 – SRR6285678). The bioinformatics analysis was deposited in a GitHub repository and permanently archived (https://doi.org/10.5281/zenodo.1188710). R scripts and corresponding data for this study have been deposited in a separate GitHub repository, which has been permanently archived (https://doi.org/10.5281/zenodo.3365703).

## Author Contributions

B.H, L.R.H, L.L.H and N.B conceived and designed the study. H.C.R and N.B contributed samples for processing. L.R.H performed laboratory work and analysed the data. I.P.A and E.L offered advice on and supervised sequencing. C.H assisted with bioinformatics analysis. P.B and S.P contributed datasets for analysis. L.R.H wrote the manuscript, which all authors revised.

