## Supporting Information for "Generating and testing ecological hypotheses at the pondscape with environmental DNA metabarcoding: a case study on a threatened amphibian"

### Contents


Appendix 1: Materials and methods 3
1.1. Samples 3
1.2 DNA reference database construction 3
1.3 Primer validation 5
1.4 eDNA metabarcoding 5
1.5 Data analysis 8
1.5.1 Preliminary analysis to identify species associations 8
1.5.2 Biotic and abiotic determinants of *T. cristatus* occupancy 9
Appendix 2: Results 11
Primer validation 11
Preliminary analysis to identify species associations 11
Appendix 3: Tables 13
Table S1 13
Table S2 15
Table S3 18
Table S4 20
Table S5 21
Table S6 22
Appendix 4: Figures 25
Figure S1 25
Figure S2 26
Figure S3 27
Figure S4 28
Figure S5 29
References 30

### Appendix 1: Materials and methods

#### 1.1 Samples

In accordance with the eDNA sampling methodology outlined by Biggs et al. (2015), 20 x 30 ml water samples were collected at equidistant intervals around the pond margin and pooled in a sterile 1 L Whirl-Pak® stand-up bag, which was shaken to provide a single homogenised sample from each pond. Six 15 ml subsamples were taken from the mixed sample using a sterile plastic pipette (25 ml) and added to sample tubes, containing 33.5 ml absolute ethanol and 1.5 ml sodium acetate 3 M (pH 5.2), for ethanol precipitation. Subsamples were then sent to Fera Science Ltd (Natural England) and ADAS (private contracts) for eDNA analysis according to laboratory protocols established by Biggs et al. (2015). Subsamples were centrifuged at 14,000 x g for 30 minutes at 6 $^{o}$C and the supernatant discarded. Subsamples were then pooled during the first step of DNA extraction with the DNeasy Blood & Tissue Kit (Qiagen®, Hilden, Germany), where 360 μl of ATL buffer was added to the first tube, vortexed, and the supernatant transferred to the second tube. This process was repeated for all six tubes. The supernatant in the sixth tube, containing concentrated DNA from all six subsamples, was transferred in a 2 ml tube and extraction continued following manufacturer's instructions to produce one eDNA sample per pond. In 2015, samples were analysed for the great crested newt (*Triturus cristatus*) using real-time quantitative PCR (qPCR) and published primers (Thomsen et al., 2012).

#### 1.2 DNA reference database construction

For freshwater fish, we used a previously created database comprising 67 fish species, which includes all known native and non-native species in the UK and the PCR positive control *Rhamphochromis esox*, a species of cichlid from Lake Malawi (Hänfling et al., 2016). For all remaining vertebrate species recorded in the UK, we used the reference databases constructed by Harper et al. (2018) in October 2016 with the ReproPhylo environment (Szitenberg, John, Blaxter, & Lunt, 2015) in a Jupyter notebook (Kluyver et al., 2016). Harper et al. (2018) deposited the Jupyter notebooks detailing the processing steps for each vertebrate group in a dedicated GitHub repository (https://github.com/HullUni-bioinformatics/Harper_et_al_2018), which has been permanently archived (https://doi.org/10.5281/zenodo.1188710).

Briefly, Harper et al. (2018) constructed species lists containing the binomial nomenclature of UK vertebrate species using the Natural History Museum UK Species Database. A BioPython script performed a GenBank search based on the species lists and downloaded all available mitochondrial 12S ribosomal RNA (rRNA) sequences for specified species. Where there were no records on GenBank for a UK species, the database was supplemented with downloaded sequences belonging to sister species in the same genus. Species that had no 12S rRNA records on GenBank are provided in Table S2. Redundant sequences were removed by clustering at 100% similarity using vsearch v1.1 (Rognes, Flouri, Nichols, Quince, & Mahé, 2016). Due to high proportion of partial 12S rRNA records on GenBank for the majority of UK species, only sequences longer than 500 bp were processed initially to increase alignment robustness to large gaps. Sequences were aligned using MUSCLE (Edgar, 2004). Short sequences can cause problems in global paired alignments where the alignment algorithm attempts to align them to longer sequences. Short 12S rRNA sequences (<500 bp) were later incorporated into the existing long 12S rRNA alignment using the hmmer v3 program suite (HMMER development team, 2016) to construct a Hidden Markov Model alignment containing sequences of all lengths. Alignments were trimmed using trimAl (Capella-Gutiérrez, Silla-Martínez, & Gabaldón, 2009). Maximum likelihood trees were inferred with RAxML 8.0.2 (Stamatakis, 2006) using the GTR+gamma model of substitutions. The complete alignments were then processed using SATIVA (Kozlov, Zhang, Yilmaz, Glöckner, & Stamatakis, 2016) for automated identification of 'mislabelled' sequences which could cause conflict in downstream analyses. Putatively mislabelled sequences were removed and process of alignment and phylogenetic tree construction repeated for manual investigation of sequences. The resultant databases (i.e. curated non-redundant reference databases) contained: 198 amphibian sequences from 20/21 species, 112 reptile sequences from 19/20 species, 272 fish sequences from 60/62 species, 940 mammal sequences from 95/112 species, and 622 bird sequences from 347/621 species. Databases for each vertebrate group were concatenated and the combined vertebrate database used for *in silico* validation of primers.

Harper et al. (2018) supplemented the amphibian database with Sanger sequences obtained from tissue of *T. cristatus*, smooth newt (*Lissotriton vulgaris*), Alpine newt (*Ichthyosaura alpestris*), common toad (*Bufo bufo*), which were supplied by University of Kent under licence from Natural England, and common frog (*Rana temporaria*), supplied by University of Glasgow. Reference sequences of the entire 12S rRNA region were generated by three sets of novel primers:

| | | |
|---|---|---|
| ***T. cristatus* (61 °C)**: | Newt_F1 | 5'-GCACTGAAAATGCTAAGACAGA-3' |
| | Newt_R6 | 5'-CAGGTATTTTCTCGGTGTAAGCA-3' |
| **Newts (59 °C):** | Newt_F2 | 5'-GCACTGAAAATGCTAAGACAG-3' |
| | Newt_R1 | 5'-TCTCGGTGTAAGCAAGATGC-3' |
| **Anura (57 °C):** | AnuraShort_F2 | 5'-TCCACTGGTCTTAGGAGCCA-3' |
| | AnuraShort_R1 | 5'-ACCATGTTACGACTTGCCTC-3' |

Harper et al. (2018) designed primers from an alignment of tRNA, 12S rRNA and 16S rRNA regions in UK Caudata and Anura species. PCR reactions were performed in 25 µl volumes containing: 12.5 µl of MyTaq™ Red Mix (Bioline Reagents Limited, London, UK), 1 µl (final concentration - 0.04 µM) of forward and reverse primer (Integrated DNA Technologies,

Belgium), 8.5µl of molecular grade sterile water (Fisher Scientific UK Ltd, Loughborough, UK) and 2 µl DNA template. PCRs were performed on an Applied Biosystems® Veriti Thermal Cycler (Fisher Scientific UK Ltd, Loughborough, UK) with the following profile: 95 °C for 3 min, 35 cycles of 95 °C for 30 sec, x °C (see temperatures above) for 60 sec and 72 °C for 90 sec, followed by a final elongation step at 72 °C for 10 min. Purified PCR products were Sanger sequenced directly (Macrogen Europe, Amsterdam, Netherlands) in both directions using the PCR primers. Sequences were edited using CodonCode Aligner (CodonCode Corporation, Centerville, MA, USA). The complete reference database compiled in GenBank format was deposited by Harper et al. (2018) in a GitHub repository for their study (https://github.com/HullUni-bioinformatics/Harper_et_al_2018), and was permanently archived (https://doi.org/10.5281/zenodo.1188710).

#### 1.3 Primer validation

Vertebrate DNA from eDNA samples was amplified with published 12S rRNA primers 12S-V5-F (5'-ACTGGGATTAGATACCCC-3') and 12S-V5-R (5'-TAGAACAGGCTCCTCTAG-3') (Riaz et al., 2011). Harper et al. (2018) validated the primers *in silico* using ecoPCR software (Ficetola et al., 2010) against their custom, phylogenetically curated reference database for UK vertebrates. Parameters were set to allow a fragment size of 50-250 bp and maximum of three mismatches between the primer pair and each sequence in the reference database. Primers were previously validated *in vitro* for UK fish communities by Hänfling et al. (2016) and by Harper et al. (2018) against tissue DNA extracted from UK amphibian species: *T. cristatus*, *L. vulgaris*, palmate newt (*Lissotrition helveticus*), *I. alpestris*, *R. temporaria* and *B. bufo*. Harper et al. (2018) performed a dilution series ($10^0$ to $10^{-8}$) for DNA (standardised to 5 ng/µl) from each amphibian species to identify the Limit of Detection for each species. Molecular grade sterile water (Fisher Scientific UK Ltd, Loughborough, UK) substituted template DNA for the PCR negative control.

#### 1.4 eDNA metabarcoding

A two-step PCR protocol was performed on eDNA samples by Harper et al. (2018). Dedicated rooms were available for pre-PCR and post-PCR processes. Pre-PCR processes were performed in a dedicated eDNA laboratory, with separate rooms for filtration, DNA extraction and PCR preparation of sensitive environmental samples. PCR reactions were set up in an ultraviolet and bleach sterilised laminar flow hood. Eight-strip PCR tubes with individually attached lids were used instead of 96-well plates to minimise cross-contamination risk between samples (Port et al., 2016) and reactions prepared for the second sequencing run were sealed with mineral oil as an additional measure against PCR contamination. For the first PCR, three replicates were performed for each sample to combat PCR stochasticity. Alternating PCR positive and negative controls were included on each PCR strip (six positive and negative controls on each 96-well plate), to screen for sources of potential contamination. The DNA used for the PCR positive control was *R. esox,* as occurrence in UK ponds is extremely rare or non-existent. The negative control substituted molecular grade sterile water (Fisher Scientific UK Ltd, Loughborough, UK) for template DNA.

During the first PCR, the target region was amplified using the primers described above, including adapters (Illumina, 2011). First PCR reactions were performed in a final volume of 21.1 µl, using 2 µl of DNA extract as a template. The amplification mixture contained 10.5 µl of MyTaq™ HS Red Mix (Bioline Reagents Limited, London, UK), 1.05 µl (final concentration - 0.5 µM) of forward and reverse primer (Integrated DNA Technologies, Belgium) and 6.5 µl of molecular grade sterile water (Fisher Scientific UK Ltd, Loughborough, UK). PCR was performed on an Applied Biosystems® Veriti Thermal Cycler (Fisher Scientific UK Ltd, Loughborough, UK) and PCR conditions consisted of an incubation step at 98 °C for 5 min, followed by 35 cycles of denaturation at 98 °C for 15 s, annealing at 56 °C for 20 s, and extension at 72 °C for 30 s, with final extension at 72 °C for 10 min. PCR products were stored at 4 °C until fragment size was verified by visualising 5 µl of selected PCR products on 2% agarose gels (100 ml 0.5x TBE buffer, 2 g agarose powder). Gels were then stained with ethidium bromide and imaged using Image Lab Software (Bio-Rad Laboratories Ltd, Watford, UK). A PCR product was deemed positive where there was an amplification band on the gel that was of the expected size (200-300 bp). PCR replicates for each sample were pooled in preparation for the addition of Illumina indexes in the second PCR, which resulted in 63.3 µl of PCR product for each sample. PCR positive and negative controls were not pooled to allow individual purification and sequencing of all 228 PCR controls. All PCR products (30 µl samples and 15 µl PCR controls) were then purified to remove excess primer using E.Z.N.A.® Cycle Pure V-Spin Clean-Up Kits (Omega Bio-tek, GA, USA) following manufacturers protocol. Eluted DNA was stored at -20 °C until the second PCR could be performed.

In the second PCR, Multiplex Identification (MID) tags (unique 8-nucleotide sequences) and Illumina MiSeq adapter sequences were bound to the amplified product. These tags were included in the forward and reverse primers resulting in indexed primers for second PCR (O'Donnell, Kelly, Lowell, & Port, 2016). For each second PCR plate, 96 unique tag combinations were created by combining eight unique forward tags with 12 unique reverse tags or vice versa (Kitson et al., 2019). A total of 384 unique tag combinations were achieved, allowing samples to be distinguished during bioinformatics analysis. Second PCR reactions were performed in eight-strip PCR tubes with individually attached lids in a final volume of 21.1 µl, using 2 µl of purified DNA from the first PCR product as a template. The amplification mixture contained 10.5 µl of MyTaq™ HS Red Mix (Bioline Reagents Limited, London, UK), 2.1 µl (final concentration - 0.5 µM) of tagged primer mix (Integrated DNA Technologies, Belgium) and 6.5 µl of molecular grade sterile water (Fisher Scientific UK Ltd, Loughborough, UK). PCR was performed on an Applied

Biosystems® Veriti Thermal Cycler (Fisher Scientific UK Ltd, Loughborough, UK) with the following profile: denaturation at 95 °C for 3 min, followed by 12 cycles of annealing at 98 °C for 20 s and extension at 72 °C for 30 s, with final extension at 72 °C for 5 min. PCR products were stored at 4 °C before they were all visualised on 2% agarose gels (100 ml 0.5x TBE buffer, 2 g agarose powder) using 5 µl PCR product. Gels were then stained with ethidium bromide and imaged using Image Lab Software (Bio-Rad Laboratories Ltd, Watford, UK). Again, PCR products were deemed positive where there was an amplification band on the gel that was of the expected size (200-300 bp).

All remaining library preparation was conducted at Fera Science Ltd. PCR products were transferred to a new 96-well PCR plate for individual purification with AMPure® XP beads (Beckman Coulter (UK) Ltd, High Wycombe, UK) and an invitrogen® magnetic stand (Fisher Scientific UK Ltd, Loughborough, UK). The Illumina PCR clean-up protocol was adapted to use 18.6 µl AMPure® XP beads (1.2x PCR product) to 15-16 µl PCR product. Illumina protocol was then followed until the beads were resuspended in 15 µl molecular grade water and incubated at room temperature for 5 minutes. The supernatant without beads in each well were not transferred to a new plate due to low volumes of purified product. Further pipetting may have resulted in loss of DNA. Each plate was sealed and stored at 4 °C until quality assurance. An Invitrogen™ Quant-IT™ PicoGreen™ dsDNA Assay (Fisher Scientific UK Ltd, Loughborough, UK) was conducted for all samples on a Fluoroskan™ Microplate Fluorometer (Life Technologies Ltd, Paisley, UK). Samples were then normalised and pooled to create 4 nM pooled libraries before quantification using an Invitrogen™ Qubit™ dsDNA HS Assay Kit (Fisher Scientific UK Ltd, Loughborough, UK). Both libraries passed quality assurance with concentrations of 2.62 ng/µl and 4.14 ng/µl respectively. An Agilent 4200 Tapestation System (Agilent Technologies, Santa Clara, CA, United States) was then used to check and compare size of the pooled libraries to selected samples. The pooled libraries were 272 bp and 299 bp (expected 286 bp) with samples in the same range. Equimolar libraries (4 nM) were then created using tapestation trace size estimates and Qubit concentrations. Libraries were run at 12 pM concentration on an Illumina MiSeq using 2 x 300 bp V3 chemistry (Illumina Inc., San Diego, CA, USA). Both libraries included a 10% PhiX DNA spike-in control to improve clustering during initial sequencing.

Illumina data was converted from raw sequences to taxonomic assignment using a custom pipeline for reproducible analysis of metabarcoding data: metaBEAT (metaBarcoding and eDNA Analysis Tool) v0.97.7 (https://github.com/HullUni-bioinformatics/metaBEAT). Bioinformatic analysis using metaBEAT largely followed the workflow outlined by Hänfling et al. (2016) for sample processing and taxonomic assignment of sequenced eDNA samples from Windermere. Adaptations to this workflow are described (see also Harper et al. 2018): raw reads were quality trimmed using Trimmomatic v0.32 (Bolger, Lohse, & Usadel, 2014), both from the read ends (minimum per base phred score Q30), as well as across sliding windows (window size 5bp; minimum average phred score Q30). Reads were clipped to a maximum length of 110 bp and reads shorter than 90 bp after quality trimming were discarded. To reliably exclude adapters and PCR primers, the first 25 bp of all remaining reads were also removed. Sequence pairs were merged into single high quality reads using FLASH v1.2.11 (Magoč & Salzberg, 2011), if a minimum of 10 bp overlap with a maximum of 10% mismatch was detected between pairs. For reads that were not successfully merged, only forward reads were kept. To reflect our expectations with respect to fragment size, a final length filter was applied and only sequences of length 80-120 bp were retained. These were screened for chimeric sequences against our custom reference database using the uchime algorithm (Edgar, Haas, Clemente, Quince, & Knight, 2011), as implemented in vsearch v1.1 (Rognes et al., 2016). Redundant sequences were removed by clustering at 97% identity ('--cluster_fast' option) in vsearch v1.1 (Rognes et al., 2016). Clusters represented by less than five sequences were considered sequencing error and omitted from further analyses. Non-redundant sets of query sequences were then compared against our custom reference database using BLAST (Zhang, Schwartz, Wagner, & Miller, 2000). For any query matching with at least 98% identity to a reference sequence across more than 80% of its length, putative taxonomic identity was assigned using a lowest common ancestor (LCA) approach based on the top 10% BLAST matches. Sequences that could not be assigned (non-target sequences) were subjected to a separate BLAST search against the complete NCBI nucleotide (nt) database at 98% identity to determine the source via LCA as described above. To ensure reproducibility of analyses, the bioinformatic analysis was archived (https://doi.org/10.5281/zenodo.1188710) by Harper et al. (2018).

#### 1.5 Data Analysis

##### 1.5.1 Preliminary analysis to identify species associations

Vertebrate species associations were investigated using the community data generated by eDNA metabarcoding with the method of Veech (2013) implemented in the R package cooccur v1.3 (Griffith, Veech & Marsh, 2016). This is a probabilistic model which measures species co-occurrence (presence-absence) as the number of sampling sites where two species co-occur. The observed co-occurrence of a given dataset is compared to the expected co-occurrence. Expected co-occurrence is determined by the probabilities of each species' occurrence multiplied by the number of sampling sites. Effect sizes were also computed for species pairs to examine species associations regardless of statistical significance. These are equivalent to the difference between expected and observed frequency of co-occurrence. The values are then standardized by dividing these differences by sample size. In standardized form, these values are bounded from -1 to 1, with positive values indicating positive associations and negative values indicating negative associations.

##### 1.5.2 Biotic and abiotic determinants of *T. cristatus* occupancy

Collinearity and spatial autocorrelation within the dataset were investigated before the most appropriate regression model was determined. Collinearity between explanatory variables was assessed using a Spearman's rank pairwise correlation matrix. Collinearity was observed between pond circumference, pond length, pond width, and pond area. Pond area encompasses length and width thus taking the same measurements and accounting for the same variance in the data as these variables. Therefore, pond circumference, pond length, and pond width were removed from the dataset so as remaining variables were not highly correlated (Zuur et al., 2009). Shading (percentage of total pond margin shaded) and terrestrial overhang (percentage of pond overhung by trees and shrubs) were also collinear. Terrestrial overhang accounts for shading of the entire pond whereas shading considers the pond margin. Shading is also a known driver of pond biodiversity (Sayer et al., 2012), thus shading was retained as an explanatory variable. Habitat Suitability Index (HSI) score was not collinear with other variables but many of the variables are also used as indices to calculate HSI score. To prevent HSI score masking variation caused by these individual variables, we analysed HSI score in a separate model. After collinear variables were removed, variance inflation factors (VIFs) of remaining variables were calculated using the package car v2.1-6 (Fox & Weisberg, 2011) to identify remnant multicollinearity. Multicollinearity (VIF < 3) (Zuur *et al.*, 2009) was not present between the candidate variables.

A large number of biotic (presence of amphibians, waterfowl and fish, *B. bufo* detection, *L. vulgaris* detection, common carp [*Cyprinus carpio*] detection, ninespine stickleback [*Pungitius pungitius*] detection, three-spined stickleback [*Gasterosteus aculeatus*] detection, common coot [*Fulica atra*] detection, common moorhen [*Gallinula chloropus*] detection) and abiotic (max. depth, pond area, pond density, shading, macrophyte cover, permanence, water quality, pond substrate, inflow, outflow, pollution, woodland, rough grass, scrub/hedge, ruderals) explanatory variables remained. The relative importance of these in explaining *T. cristatus* detection was inferred using a classification tree within the package rpart v4.1-13 (Therneau, Atkinson & Ripley, 2014). The classification tree suggested the most important explanatory variables of *T. cristatus* detection were: *L. vulgaris* detection, fish presence, *B. bufo* detection, amphibian presence, pond area, *G. chloropus* detection, pond substrate, water quality, pond density, woodland, permanence, max. depth, outflow, inflow, scrub/hedge, percentage of macrophyte cover, percentage of shading, ruderals, and waterfowl presence. *L. vulgaris*, *B. bufo*, and *G. chloropus* were also identified as having significant associations with *T. cristatus* by the preliminary cooccur analysis. A pruning diagram was applied to the data to cross-validate the classification tree and remove unimportant explanatory variables. A tree of 4 was optimal according to the pruning diagram, indicating that 4 explanatory variables should be retained for statistical analysis. Although not identified by the classification tree, we decided to include presence of *C. carpio*, *G. aculeatus* and *P. pungitius* in models as these fish directly predate *T.*

*cristatus*, and *F. atra* presence as a common waterfowl species that prefers similar habitat to *T. cristatus* and may compete for resources. Many variables occurred more than once in the classification tree, indicative of weak non-linear relationships with the response variable. Generalized Additive Models (GAMs) were performed to assess non-linearity but several explanatory variables were in fact linear, i.e. estimated one degree of freedom for smoother (Zuur et al., 2009).

The ponds in this study had restricted spatial distribution and were nested within three UK counties (Fig. S1) thus spatial autocorrelation may be present. This phenomena is common in ecological studies of species presence-absence as sites located within an animal's ranging capability are likely to be inhabited (Zuur et al., 2009). *T. cristatus* individuals can migrate distances of 1-2 km to new ponds (Edgar & Bird, 2006; Haubrock & Altrichter, 2016), thus *T. cristatus* are likely to occupy ponds that are closely located to one another in a given county. Spline correlograms - graphical representations of spatial correlation between locations at a range of lag distances that are smoothed using a spline function (Bjørnstad, 2017) - were constructed using the package ncf v1.1-7 to examine spatial autocorrelation between ponds. Spline correlograms of the Pearson residuals of the raw data, a binomial Generalized Linear Model (GLM), and a binomial Generalized Linear Mixed-effects Model (GLMM) were compared. GLMMs can account for dependencies within sites, handled with the introduction of random effects (Zuur et al., 2009). As ponds were clustered within three counties, county was treated as a random effect. The GLMM successfully accounted for spatial dependencies between ponds based on the spline correlogram of the Pearson residuals. After identifying a suitable set of explanatory variables and modelling framework, we constructed separate binomial GLMMs with the logit link function for biotic and abiotic explanatory variables. For each GLMM, we used an information-theoretic approach using Akaike Information Criterion (AIC) to determine the most parsimonious approximating model to make predictions (Akaike, 1973).

### Appendix 2: Results

#### 2.1 Primer validation

The *in silico* analysis performed by Harper et al. (2018) confirmed taxonomic coverage (59.0% of target vertebrate species amplified) and resolution of the 12S rRNA primers. A wide range of UK vertebrate taxa were amplified, with fragment length ranging from 90-114 bp. The primers amplified 16/21 amphibian species, including *T. cristatus*. *L. helveticus*, Italian crested newt (*Triturus carnifex*), brown cave salamander (*Hydromantes genei*), marsh frog (*Pelophylax esculentus*) and agile frog (*Rana dalmatina*) were not amplified *in silico*. Harper et al. (2018) manually aligned all sequences from these species to the primers using the alignment viewer and editor AliView (Larsson, 2014), confirming potential for amplification. The primers amplified 47/67 fish species, including the threatened European eel (*Anguilla anguilla*), but amplification of UK freshwater fish assemblages was confirmed *in vitro* by Hänfling et al. (2016). The primers amplified 14/20 reptile species including slow worm (*Anguis fragilis*) and common lizard (*Zootoca vivipara*). Reference sequences were not available for one species and a further five species were not amplified. Primers were only validated for 282/621 bird species (including common waterfowl species). There were no 12S rRNA data available for 243/621 bird species and a further 96 species were not amplified. Similarly, no reference data were available for nine mammal species (bats and marine mammals) and a further 15 species were not amplified. Only 88/112 mammal species were validated. Several marine mammal species were not amplified but would not be found in freshwater ponds. However, priority species for freshwater management, such as water vole *Arvicola amphibius*, were not amplified alongside other species of bat, vole and shrew that may frequent ponds. Harper et al. (2018) observed bands by agarose gel electrophoresis for all amphibian tissue they tested, including *L. helveticus* which was not amplified *in silico*, and no bands were observed in NTCs. The Limit of Detection was variable for each species: *T. cristatus*, *L. helveticus*, *R. temporaria* and *B. bufo* were not amplified below $5 \times 10^{-4}$ ng/µl, whereas *I. alpestris* was not amplified below $5 \times 10^{-3}$ ng/µl and *L. vulgaris* below $5 \times 10^{-5}$ ng/µl. Due to sheer number of and legislation surrounding many UK amphibian, reptile, bird and mammal species, *in vitro* testing for all target taxa is unfeasible and metabarcoding proceeded on the basis of *in silico* amplification.

#### 2.2 Preliminary analysis to identify species associations

The cooccur analysis revealed that 1083 (81.67%) of 1326 species pair combinations were removed from the analysis because expected co-occurrence was less than one, leaving 243 pairs for analysis. The pairwise combinations revealed 17 negative and 31 positive significant co-occurrence patterns. The remaining co-occurrence patterns were random thus the observed presence-absence data did not significantly deviate from the expected presence-absence data. No pairs were unclassifiable indicative of sufficient statistical power to analyse all pairs. A pairing profile was constructed to understand each species' individual contribution to the positive and negative species associations. Interactions were clustered in a few species rather than being evenly distributed. When observed and expected co-occurrence was examined, some species pairs deviated from the expected co-occurrence. A minority of species pairs exhibited fewer than expected co-occurrences but these pairs were largely clustered towards having low expected co-occurrence.

#### Appendix 3: Tables

**Table S1.** Summary of environmental metadata on pond characteristics and surrounding terrestrial habitat collected by environmental consultants contracted for Natural England's Great Crested Newt Evidence Enhancement Programme.

| Variable | Description | Unit/categories |
|---|---|---|
| Maximum depth | Depth of pond | m |
| Circumference | Pond circumference | m |
| Width | Pond width | m |
| Length | Pond length | m |
| Area | Pond area | $m^2$ |
| Density | Pond density | Number of ponds per $km^2$ |
| Terrestrial overhang | Percentage of pond overhung by trees and shrubs | % |
| Shading | Percentage of total pond margin shaded to at least 1 m from the shore | % |
| Macrophyte cover | Percentage of pond surface occupied by macrophytes | % |
| Habitat Suitability Index (HSI) | Score calculated from aforementioned variables which indicates habitat quality for crested newt (0 = poor, 1 = excellent) | Decimal |
| Habitat Suitability Index (HSI) band | Categorical classification of HSI score | Poor/below average/average/good |
| Pond permanence | Pond permanence | Dries annually/rarely dries/sometimes dries/ never dries |
| Water quality | Subjective assessment based on | Bad/poor/moderate/go |

| Variable | Description | Unit/categories |
| --- | --- | --- |
| | invertebrate diversity, presence of submerged vegetation, and knowledge of water inputs to pond. | od/excellent |
| Pond substrate | Type of substrate | Not known/rock/clay/concrete/sand, gravel, pebbles/lined/peat-organic |
| Inflow | Water inputs to pond | Absent/present |
| Outflow | Water leaving pond | Absent/present |
| Pollution | Rubbish or other signs of pollution | Absent/present |
| Other amphibians | Presence of amphibian species other than crested newt | Absent/present |
| Fish | Presence of any fish species | Absent/possible/minor/major |
| Waterfowl | Presence of any waterfowl species | Absent/minor/major |
| Woodland | Terrestrial habitat: woodland | None/some/important |
| Rough grass | Terrestrial habitat: rough grass | None/some/important |
| Scrub/hedge | Terrestrial habitat: scrub/hedge | None/some/important |
| Ruderals | Terrestrial habitat: ruderals | None/some/important |
| Terrestrial other | Other good quality terrestrial habitat that does not conform to aforementioned habitat types | None/some/important |
| Overall terrestrial habitat score | Overall quality of terrestrial habitat | None/poor/moderate/good |

**Table S2.** List of species for which no 12S rRNA records were available on GenBank**.** Only UK species which had no records for sister species within the same genus are included.

| **Common name** | **Binomial nomenclature** |
|---|---|
| North Atlantic right whale | *Eubalaena glacialis* |
| Common kingfisher | *Alcedo atthis* |
| Trumpeter finch | *Bucanetes githagineus* |
| Green heron | *Butorides virescens* |
| Greater short-toed lark | *Calandrella brachydactyla* |
| Lesser short-toed lark | *Calandrella rufescens* |
| Lapland longspur | *Calcarius lapponicus* |
| Wilson's warbler | *Cardellina pusilla* |
| Rufuous-tailed scrub robin | *Cercotrichas galactotes* |
| MacQueen's bustard | *Chlamydotis macqueenii* |
| Lark sparrow | *Chondestes grammacus* |
| White-throated dipper | *Cinclus cinclus* |
| Great spotted cuckoo | *Clamator glandarius* |
| Long-tailed duck | *Clangula hyemalis* |
| Corn crake | *Crex crex* |
| Crested lark | *Galerida cristata* |
| European storm petrel | *Hydrobates pelagicus* |
| Little gull | *Hydrocoloeus minutus* |
| White-throated robin | *Irania gutturalis* |
| Hooded merganser | *Lophodytes cucullatus* |
| European crested tit | *Lophophanes cristatus* |
| Woodlark | *Lullula arborea* |
| Siberian blue robin | *Larvivora cyane* |
| Rufous-tailed robin | *Larvivora sibilans* |
| Thrush nightingale | *Luscinia luscinia* |

| Common name | Binomial nomenclature |
|---|---|
| Common nightingale | *Luscinia megarhynchos* |
| Bluethroat | *Luscinia svecica* |
| Black scoter | *Melanitta americana* |
| Velvet scoter | *Melanitta fusca* |
| Common scoter | *Melanitta nigra* |
| Surf scoter | *Melanitta perspicillata* |
| Bimaculated lark | *Melanocorypha bimaculata* |
| Calandra lark | *Melanocorypha calandra* |
| White-winged lark | *Melanocorypha leucoptera* |
| Black lark | *Melanocorypha yeltoniensis* |
| Song sparrow | *Melospiza melodia* |
| Black-and-white warbler | *Mniotilta varia* |
| Common rock thrush | *Monticola saxatilis* |
| Blue rock thrush | *Monticola solitarius* |
| Wilson’s storm petrel | *Oceanites oceanicus* |
| Band-rumped storm petrel | *Oceanodroma castro* |
| Leach’s storm petrel | *Oceanodroma leucorhoa* |
| Swinhoe’s storm petrel | *Oceanodroma monorhis* |
| Tennessee warbler | *Oreothlypis peregrina* |
| Northern waterthrush | *Parkesia noveboracensis* |
| Savannah sparrow | *Passerculus sandwichensis* |
| Rosy starling | *Pastor roseus* |
| American cliff swallow | *Petrochelidon pyrrhonota* |
| Steller’s eider | *Polysticta stelleri* |
| Eurasian crag martin | *Ptyonoprogne rupestris* |
| Sand martin | *Riparia riparia* |
| Whinchat | *Saxicola rubetra* |
| African stonechat | *Saxicola torquatus* |

| **Common name** | **Binomial nomenclature** |
|---|---|
| Northern parula | *Setophaga americana* |
| Hooded warbler | *Setophaga citrina* |
| American yellow warbler | *Setophaga petechia* |
| American redstart | *Setophaga ruticilla* |
| Wallcreeper | *Tichodroma muraria* |
| Brown thrasher | *Toxostoma rufum* |
| Golden-winged warbler | *Vermivora chrysoptera* |

**Table S3.** List of species detected in PCR positive controls by eDNA metabarcoding and corresponding taxon-specific false positive sequence threshold applied.

| Binomial name | Common name | Taxon-specific false positive sequence threshold |
|---|---|---|
| Actinopteri | Actinopteri | 0.000141306 |
| *Anas* | Dabbling ducks | 0.1 |
| *Anguilla anguilla* | European eel | 0.0000939 |
| Aves | Birds | 0.133333333 |
| *Bos taurus* | Cow | 0.003542152 |
| *Bufo bufo* | Common toad | 0.066666667 |
| *Clupea harengus* | Atlantic herring | 0.000114602 |
| *Columba* | Doves | 0.000129631 |
| Columbidae | Pigeons and doves | 0.000889494 |
| Corvidae | Corvids | 0.002149471 |
| Cyprinidae | Cyprinids | 0.002535206 |
| *Cyprinus carpio* | Common carp | 0.00016315 |
| *Fulica atra* | Common coot | 0.000222549 |
| *Gallinula chloropus* | Common moorhen | 0.000178659 |
| *Gasterosteus aculeatus* | Three-spined stickleback | 0.066666667 |
| Hominidae | Great apes | 0.007432086 |
| *Homo sapiens* | Human | 0.839569452 |
| *Lissotriton vulgaris* | Smooth newt | 0.066666667 |
| Passeriformes | Passerine birds | 0.000489199 |
| Percidae | Perciform fish | 0.000734174 |
| Phasianidae | Phasianids | 0.000721061 |
| *Phoxinus phoxinus* | Common minnow | 0.001287409 |
| Primates | Primates | 0.000983552 |
| *Rana temporaria* | Common frog | 0.000596469 |
| *Rattus norvegicus* | Brown rat | 0.000466826 |

| Binomial name | Common name | Taxon-specific false positive sequence threshold |
|---|---|---|
| *Rutilus rutilus* | Common roach | 0.000291467 |
| Salmonidae | Salmonids | 0.000510068 |
| *Squalius cephalus* | European chub | 0.004080097 |
| *Sturnus vulgaris* | Common starling | 0.000138665 |
| *Sus scrofa domesticus* | Domestic pig | 0.000877385 |
| *Triturus cristatus* | Great crested newt | 0.000276159 |
| unassigned | NA | 0.266666667 |

**Table S4.** List of species removed from the dataset prior to statistical analysis.

| Common name | Binomial name | Number of eDNA samples |
|---|---|---|
| Cichlid | *Rhamphochromis esox* | 287 |
| Human | *Homo sapiens* | 7 |
| Domestic dog | *Canis lupus familiaris* | 63 |
| Horse | *Equus caballus* | 3 |
| Cow | *Bos taurus* | 177 |
| Sheep | *Ovis aries* | 42 |
| Domestic pig | *Sus scrofa domesticus* | 139 |
| Domestic cat | *Felis catus* | 16 |
| Domesticated turkey | *Meleagris gallopavo* | 11 |
| Helmeted guineafowl | *Numida meleagris* | 1 |

**Table S5.** Full a priori models used as input for stepwise backward deletion of terms, and the final models resulting from model selection.

| Model name | Full *a priori* model | Final model |
|---|---|---|
| Model 1 | *T. cristatus* ~ (1\|County) + amphibian richness + fish richness + waterfowl richness + terrestrial bird richness + mammal richness | *T. cristatus* ~ (1\|County) + amphibian richness + fish richness + waterfowl richness + terrestrial bird richness + mammal richness |
| Model 2 | *T. cristatus* ~ (1\|County) + *Lissotriton vulgaris* + *Bufo bufo* + Amphibians + Fish + Waterfowl + *Cyprinus carpio* + *Gasterosteus aculeatus* + *Pungitius pungitius* + *Gallinula chloropus* + *Fulica atra* | *T. cristatus* ~ (1\|County) + *Lissotriton vulgaris* + *Bufo bufo* + *Cyprinus carpio* + *Gasterosteus aculeatus* + *Gallinula chloropus* + Waterfowl |
| Model 3 | *T. cristatus* ~ (1\|County) + max. depth + pond density + inflow + pond area + pond substrate + outflow + macrophyte cover + water quality + permanance + shading + ruderals + scrub/hedge + woodland | *T. cristatus* ~ (1\|County) + inflow + pond area + shading |
| Model 4 | *T. cristatus* ~ (1\|County) + HSI score | *T. cristatus* ~ (1\|County) + HSI score |
| Model 5 | Vertebrate species richness ~ (1\|County) + *T. cristatus* + HSI score | Vertebrate species richness ~ (1\|County) + *T. cristatus* + HSI score |

**Table S6.** Summary of species detected by eDNA metabarcoding of freshwater ponds (*N* = 532).

| Common name | Binomial name | Number of ponds |
|---|---|---|
| European eel | *Anguilla anguilla* | 15 |
| Common barbel | *Barbus barbus* | 2 |
| Crucian carp | *Carassius carassius* | 2 |
| Common carp | *Cyprinus carpio* | 40 |
| Common minnow | *Phoxinus phoxinus* | 12 |
| Common roach | *Rutilus rutilus* | 71 |
| European chub | *Squalius cephalus* | 20 |
| Stone loach | *Barbatula barbatula* | 14 |
| Northern pike | *Esox lucius* | 17 |
| European bullhead | *Cottus gobio* | 14 |
| Three-spined stickleback | *Gasterosteus aculeatus* | 55 |
| Ninespine stickleback | *Pungitius pungitius* | 15 |
| Ruffe | *Gymnocephalus cernua* | 1 |
| Rainbow trout | *Oncorhynchus mykiss* | 3 |
| Common toad | *Bufo bufo* | 42 |
| Marsh frog | *Pelophylax ridibundus* | 1 |
| Common frog | *Rana temporaria* | 122 |
| Palmate newt | *Lissotrition helveticus* | 5 |
| Smooth newt | *Lissotriton vulgaris* | 151 |
| Great crested newt | *Triturus cristatus* | 148 |
| Dabbling ducks | *Anas* spp. | 150 |
| Eurasian oystercatcher | *Haematopus ostralegus* | 1 |
| Common buzzard | *Buteo buteo* | 4 |

| Common name | Binomial name | Number of ponds |
|---|---|---|
| Common pheasant | *Phasianus colchicus* | 25 |
| Eurasian coot | *Fulica atra* | 48 |
| Common moorhen | *Gallinula chloropus* | 211 |
| Eurasian jay | *Garrulus glandarius* | 7 |
| European goldfinch | *Carduelis carduelis* | 1 |
| Dunnock | *Prunella modularis* | 4 |
| Eurasian nuthatch | *Sitta europaea* | 1 |
| Common starling | *Sturnus vulgaris* | 4 |
| Melodius warbler | *Hippolais polyglotta* | 2 |
| Grey heron | *Ardea cinerea* | 1 |
| Great spotted woodpecker | *Dendrocopus major* | 1 |
| Green woodpecker | *Picus viridis* | 2 |
| Tawny owl | *Strix aluco* | 1 |
| Red fox | *Vulpes vulpes* | 9 |
| Eurasian otter | *Lutra lutra* | 1 |
| European badger | *Meles meles* | 7 |
| European polecat | *Mustela putorius* | 1 |
| Common pipistrelle | *Pipistrellus pipistrellus* | 1 |
| Eurasian water shrew | *Neomys fodiens* | 8 |
| Common shrew | *Sorex araneus* | 1 |
| European hare | *Lepus europaeus* | 1 |
| European rabbit | *Oryctolagus cuniculus* | 23 |
| European water vole | *Arvicola amphibius* | 16 |
| Bank vole | *Myodes glareolus* | 8 |

| Common name | Binomial name | Number of ponds |
|---|---|---|
| House mouse | *Mus musculus* | 16 |
| Brown rat | *Rattus norvegicus* | 39 |
| Grey squirrel | *Sciurus carolinensis* | 57 |
| Red deer | *Cervus elaphus* | 2 |
| Reeve's muntjac | *Muntiacus reevesi* | 3 |

### Appendix 4: Figures

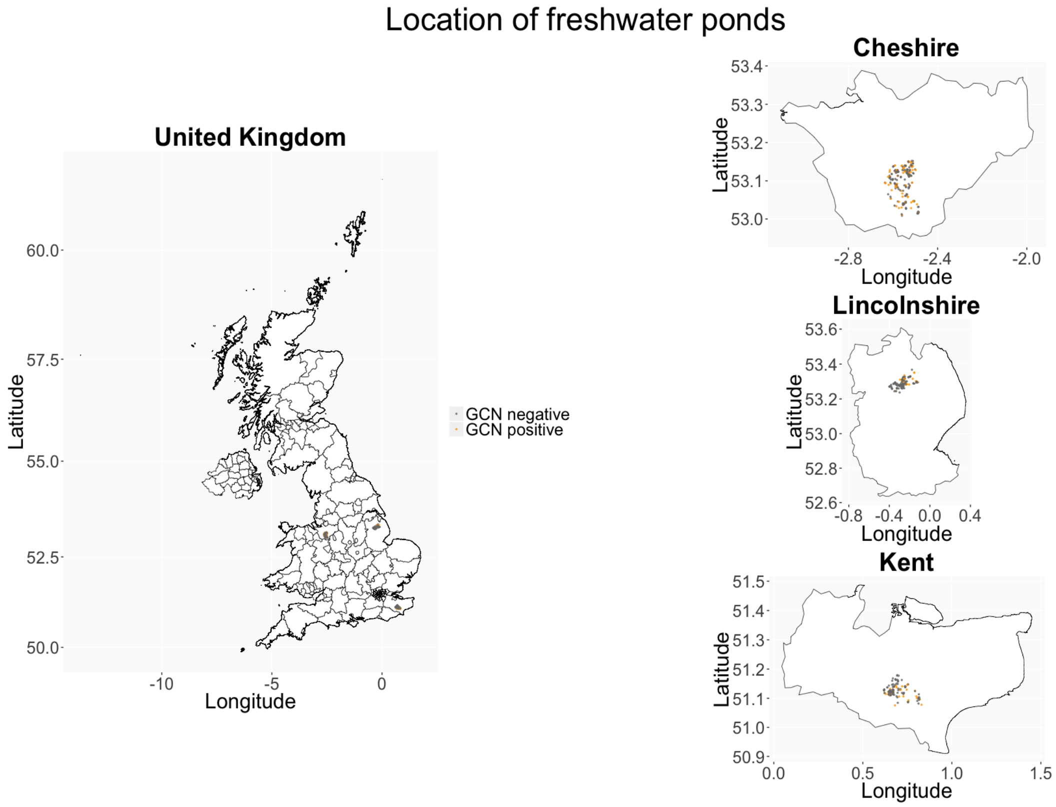


**Figure S1.** Location of ponds (*n* = 504) sampled for eDNA as part of Natural England's Great Crested Newt Evidence Enhancement Programme. Ponds that were negative or positive for *T. cristatus* (GCN) by targeted quantitative PCR are indicated by grey and orange points respectively.

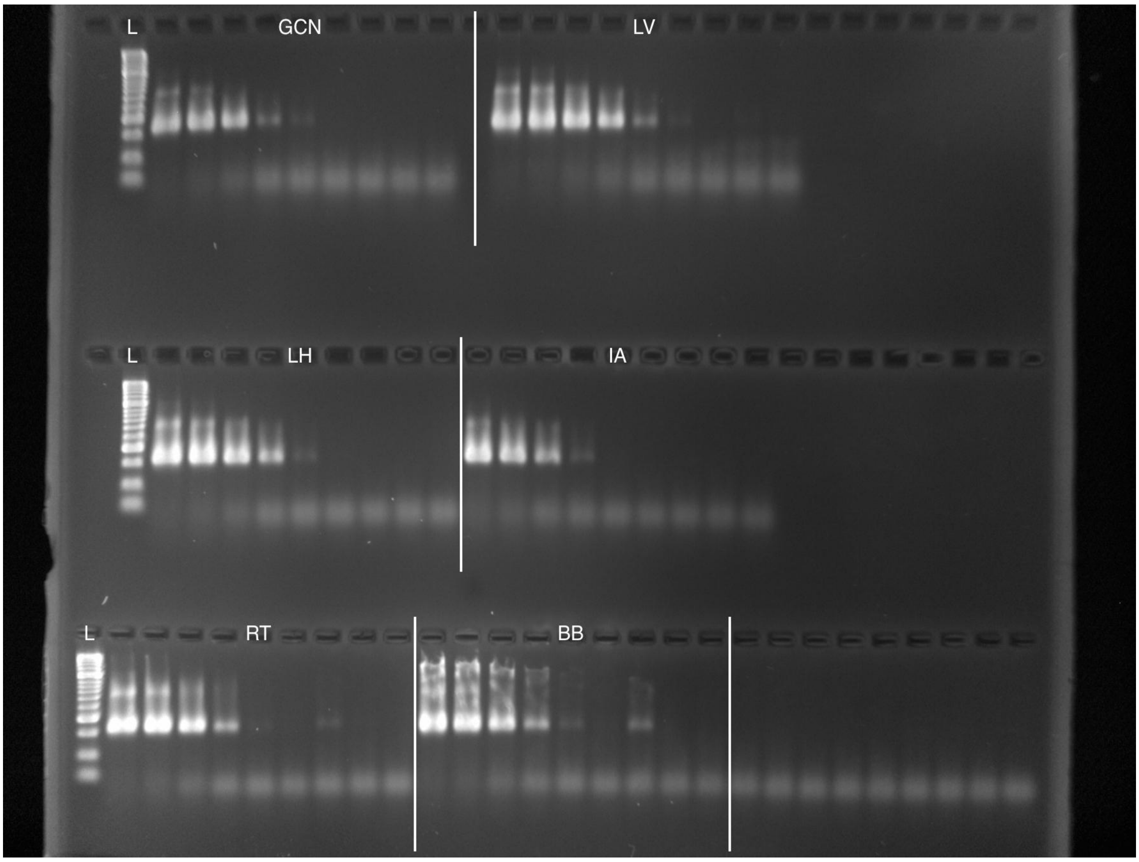


**Figure S2.** Gel image showing results of *in vitro* primer validation conducted by Harper et al. (2018)**.** All tissue DNA used for dilution series was standardised to a starting concentration of 5 ng/µl. The Limit of Detection was variable for each species: *Triturus cristatus* (GCN), *Lissotriton helveticus* (LH), *Rana temporaria* (RT) and *Bufo bufo* (BB) were not amplified below 5 x $10^{-4}$ ng/µl, whereas *Icthyosaura alpestris* (IA) was was not amplified below 5 x $10^{-3}$ ng/µl and *Lissotriton vulgaris* (LV) below 5 x $10^{-5}$ ng/µl.

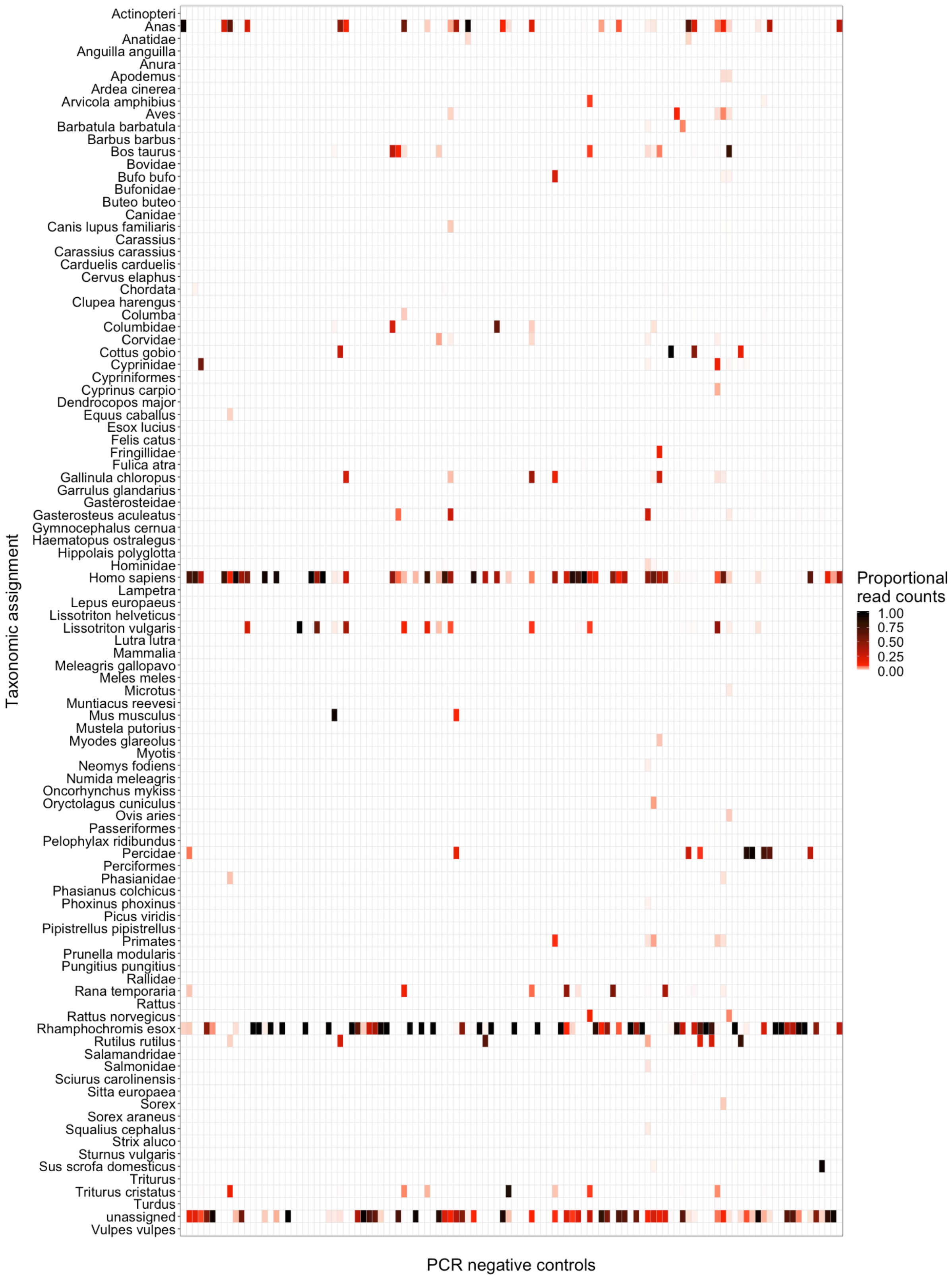


**Figure S3.** Heatmap showing the frequency of contamination in PCR negative controls. Assignments that were not detected in a PCR negative control are coloured white.

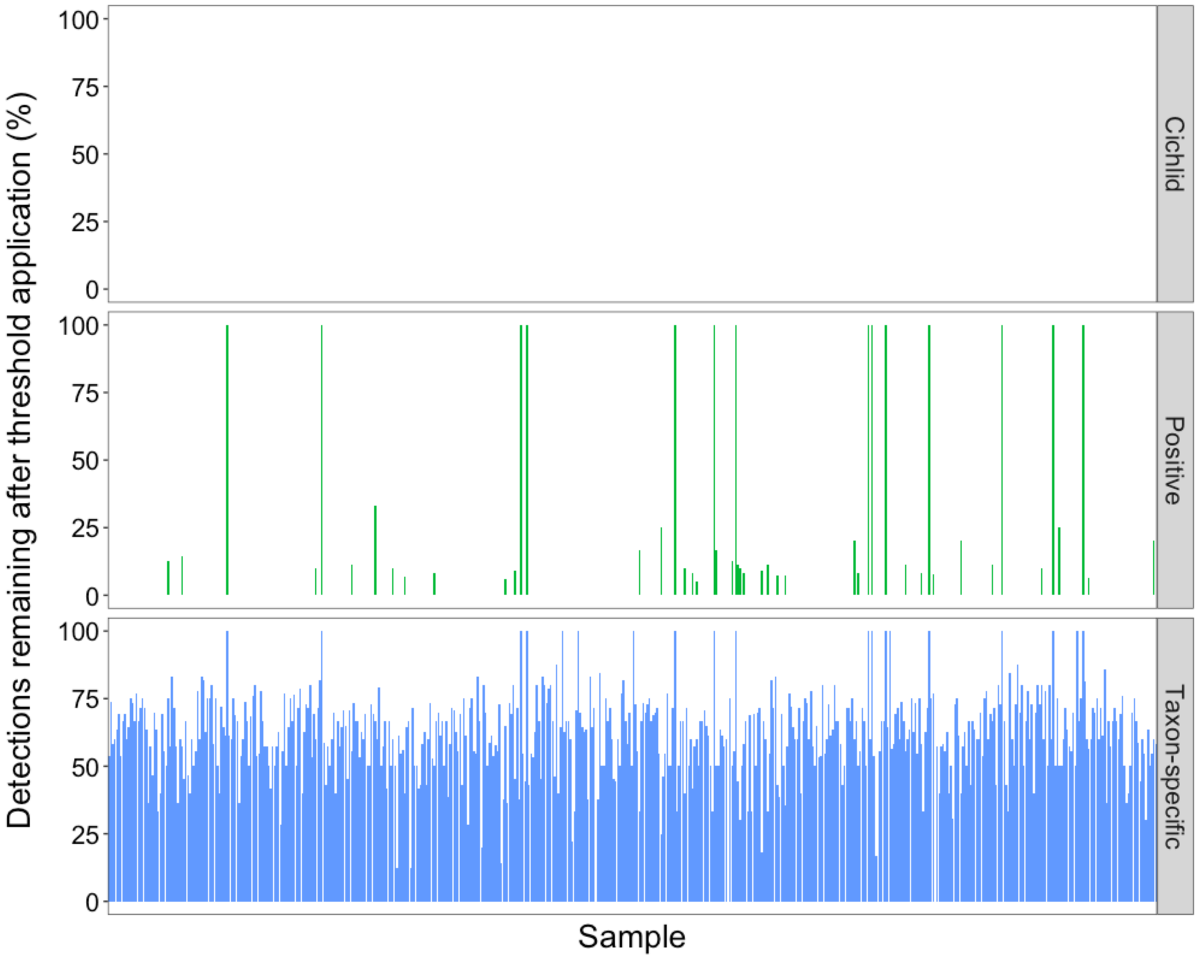


**Figure S4.** Barplot showing the impact of different false positive sequence thresholds on the proportion of taxa detected in each sample. The taxon-specific thresholds retained the most biological information, thus these were applied to the eDNA metabarcoding data for downstream analyses.

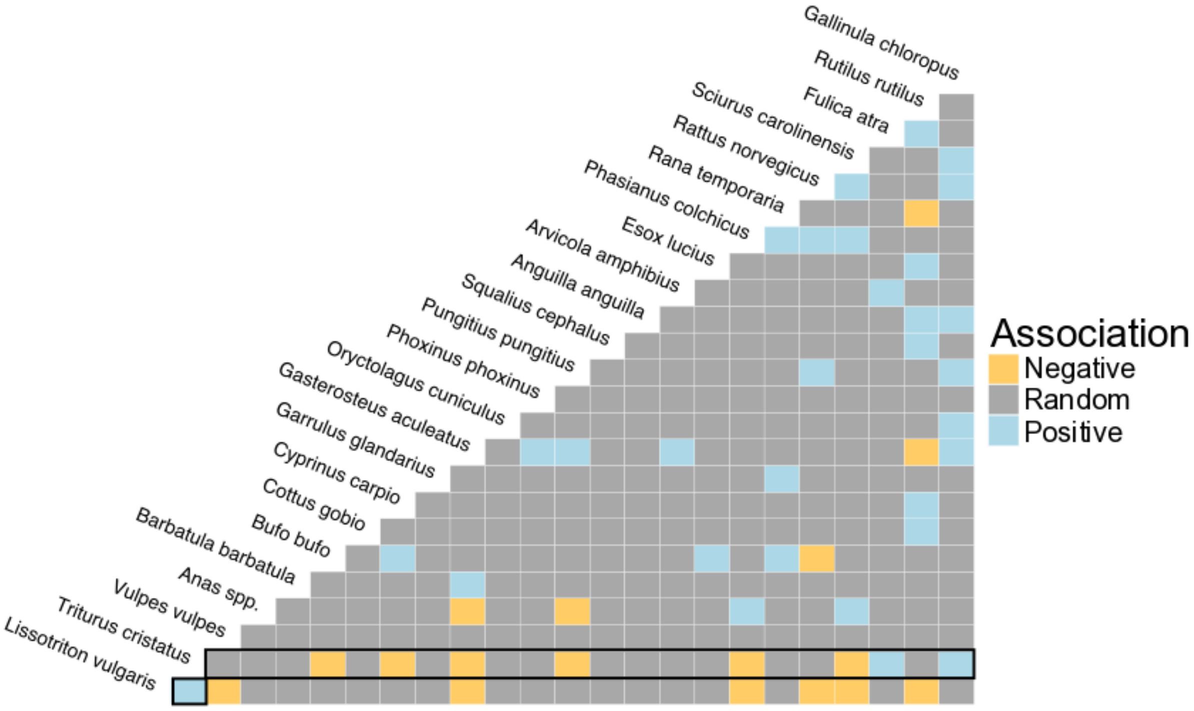


**Figure S5.** Heat map showing significant ($P < 0.05$) positive and negative species associations determined by the probabilistic co-occurrence model for the eDNA metabarcoding presence-absence data ($N$ = 532 ponds). Species names are positioned to indicate the columns and rows that represent their pairwise relationships with other species. Species are ordered by those with the most negative interactions to those with the most positive interactions (left to right). Associations relevant to *T. cristatus* are outlined in black.
